# Genomics of turions from the Greater Duckweed reveal its pathways for dormancy and reemergence strategy

**DOI:** 10.1101/2022.12.24.521731

**Authors:** Buntora Pasaribu, Kenneth Acosta, Anthony Aylward, Yuanxue Liang, Bradley W. Abramson, Kelly Colt, T. Hartwick Nolan, John Shanklin, Todd P. Michael, Eric Lam

## Abstract

- Over 15 families of aquatic plants are known to use a strategy of developmental switching upon environmental stress to produce dormant propagules called turions. However, few molecular details for turion biology have been elucidated due to the difficulties in isolating high-quality nucleic acids from this tissue. We successfully developed a new protocol to isolate high-quality transcripts and carried out RNA-seq analysis of mature turions from the Greater Duckweed *Spirodela polyrhiza*. Comparison of turion transcriptome to that of fronds, the actively growing leaf-like tissue, were carried out.
- Bioinformatic analysis of high confidence, differentially expressed transcripts between frond and mature turion tissues revealed major pathways related to stress tolerance, starch and lipid metabolism, and dormancy that are mobilized to reprogram frond meristems for turion differentiation.
- We identified the key genes that are likely to drive starch and lipid accumulation during turion formation, as well as in pathways for starch and lipid utilization upon turion germination. Comparison of genome-wide cytosine methylation levels also revealed evidence for epigenetic changes in the formation of turion tissues.
- Similarities between turions and seeds provided evidence that key regulators for seed maturation and germination have been retooled for their function in turion biology.

## Introduction

Plants and animals have evolved strategies to cope with environmental stresses, which are abrupt changes in abiotic or biotic factors, as well as patterns of cyclical changes such as seasonal variations in temperature and rainfall. As sessile organisms, plants elaborate various strategies of developmental transition into a dormant state that is often likened to hibernation in animals (Morin & Storey, 2009). For freshwater aquatic plants, many have evolved the capability to produce dormant, bud-like structures generally called turions as a stress-response and overwintering survival strategy that does not involve sexual reproduction (Adamec 2018). In the Lemnaceae family (commonly called duckweeds) of freshwater macrophytes, the species in which turions have been most well-characterized is *Spirodela polyrhiza* (aka Greater Duckweed), which has been the subject of detailed investigation for the past five decades. Specific turion-inducing factors are limiting nutrient availability (phosphate, nitrate, and sulfate) and low temperatures (Appenroth *et al*.,1989; Appenroth *et al*., 2002) both *in vitro* and under field conditions (Appenroth *et al*., 2009). Turion formation capacity, as measured by specific turion yield (STY) of different *S. polyrhiza* strains, has been quantitatively linked to local climatic conditions, such as average temperature and annual precipitation, as well as those in the growing season (Kuehdorf *et al*., 2014, Fig. S1).

Vegetative fronds of *S. polyrhiza* multiply through continuous generation of daughter fronds from meristematic tissues within two internal regions, called pockets, of the mother frond (Appenroth *et al*., 2011). Upon induction by one or more factors for turion biogenesis, the developmental program of the frond primordia in the two pockets of *S. polyrhiza* switches to produce dormant structures called turions that eventually detach from the mother fronds and sink to the bottom of the water body, where they are less likely to freeze in winter (Smart & Trewavas, 1983a; Landolt & Kandeler, 1987). The turion state could also be advantageous in areas of seasonal flooding, such as many locales in India, where low nutrient induced turions are less likely to be washed away from flooded lakes and streams when they are lying at the bottom of the water column. These dormant propagules are round shaped, dark green or brownish-green in color, and in *S. polyrhiza* have an average diameter of 1-3 mm (Fig. 1, Fig. S2). Turions have smaller cell size, lack aerenchyma which are specialized plant cells that undergo stereotypical programmed death to create internal air spaces (Smart & Trewavas, 1983b), have lower chlorophyll but higher anthocyanin content, and thicker cell walls when compared to those of normal fronds (Jacob, 1947; Appenroth et al., 2011). These characteristics help distinguish turions from other varieties of dormant states, collectively called “resting fronds,” that have been reported for a variety of duckweeds (Landolt, 1986). Like seeds, dormancy of turions is broken by cold temperatures over a span of several weeks during the overwintering process (Landolt & Kandeler, 1987). After the return of suitable conditions in the spring and together with an essential light signal (mediated via phytochrome), germination is triggered, and active growth can resume (Appenroth & Gabrys, 2001).

**Figure 1.**
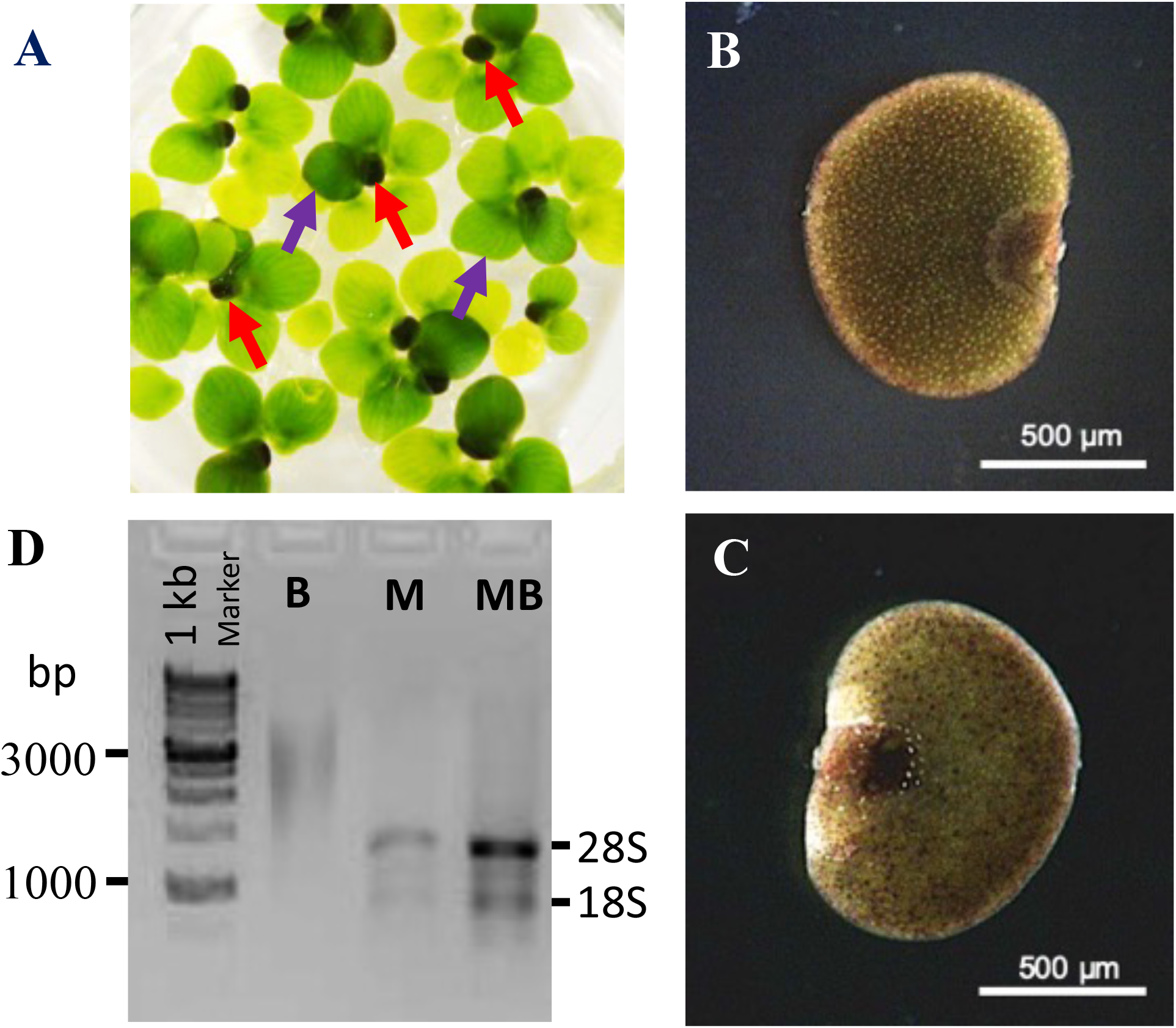
Turion of *Spirodela polyrhiza* and optimization of RNA Isolation. (A). Immature turions (dark color) are attached to the mother fronds of *Spirodela polyrhiza* under limiting phosphate conditions. (B) Ventral side, and (C) Dorsal view of mature turion. *S. polyrhiza* turion is round shaped and dark brown, indicative of high anthocyanin content. (D) Optimization of total RNA isolated from *S. polyrhiza* mature *t*urions (sp9512). B: Beads only; M: Mortar only; MB: Mortar and Beads. Turions indicated by red arrowheads. Fronds indicated by purple arrowheads.

Little is known about turion molecular biology in terms of biogenesis pathways and gene expression in this specialized developmental state. One of the major obstacles that has impeded progress is the difficulty encountered for nucleic acid isolation from turions due in part to their high starch content of more than 70% of their dry mass (Fig. S2) and high levels of tannins. To date, published molecular studies attempting to examine turion-related transcripts have been carried out with *S. polyrhiza* fronds treated with either low levels (0.25 uM) of abscisic acid (ABA) for a few hours (Smart & Flemning, 1993), or high levels (10 uM) of ABA for 3 days (Wang *et al*., 2014a). While the former study used cDNA cloning via subtractive hybridization to identify several ABA-responsive duckweed genes, and the later work applied RNA sequencing (RNA-seq) to describe several hundred transcripts that are either up- or down-regulated upon treatment with high ABA levels, their direct relevance to turion biology remains unclear since the large percentage of transcripts from frond tissues in these studies confounded the interpretation of the data. We have overcome this key hurdle to generate high quality transcriptome libraries and high molecular weight genomic DNA from turions to characterize transcript abundance and genome modification in mature turions. Our comparative analysis led us to discover heightened levels of oil production in mature turions, while mapping of 5-methycytosine abundance revealed evidence that epigenetic mechanisms may be involved in this developmental transition to modify the expression of a subset of genes as well as to provide additional genome stability during extended time of dormancy in duckweed turions.

## Material and Methods

### Duckweed material

*Spirodela polyrhiza* clones were obtained from RDSC (Rutgers Duckweed Stock Collection) at Rutgers University, New Brunswick, NJ, USA. The clones were maintained on 0.5X SH medium with 0.8% w/v agar and 0.1% sucrose with the addition of 100 mg/L cefotaxime. The clones were stored at 15°C under illumination of 40–44 μmol m^−2^ s^−1^ light. Clones selected for studies were sterilized before transfer to growth under 25°C and 150 umol m^−2^ s^−1^ light, 16hr light/8hr dark cycle.

### Turion induction

Two three-frond colonies of sp9512 and sp9509 clones from stock cultures were grown in solution containing 0.5X SH medium and 0.5% sucrose for two weeks. 200 mg fresh fronds of each clones were then transferred to 177 ml glass jar containing 50 ml of mineral salt medium: 60 μM KH_2_PO_4_, 1 mM Ca(NO_3_)_2_.4H_2_O, 8 mM KNO_3_, 5 mM H_3_BO_3_,13 μM MnCl_2_.4H_2_O, 0.4 μM Na_2_MoO_4_, 1 mM MgSO_4_.7H_2_O and 25 μM FE(III) EDTA for one week. For turion induction, 50 mg fresh frond of each clone were transferred to 177 ml glass jar containing 50 ml mineral salt medium with 2 μM (low Pi media) KH_2_PO_4_ instead of 60 μM. Mature turions are harvested when these sank to the bottom of the glass jars.

### Total RNA preparation from turion and frond tissues

Tissues for RNA preparation are collected and flash frozen in liquid nitrogen for storage at -80°C until use for nucleic acid isolation. RNA isolation from frond tissues were done with the *mir*Vana kit from Life Tech., following essentially directions from the manufacturer. For our improved method with turions, a modified CTAB method (Gambino *et al*., 2008) was used. The frozen tissues were first ground by mortar and pestle in liquid nitrogen. The fine powder was transferred to bead beating tubes with silica beads, then filled with extraction buffer, freeze in liquid nitrogen for 30 sec., and bead beating at 4°C for 15 min. before extraction of the nucleic acid. Detailed protocols for these procedures can be found in the Supporting Information file.

### RNA library preparation and sequencing

RNA samples were shipped and sequenced separately at either BGI Genomics (Hong Kong, China) or Novogene Co., Ltd (Beijing, China), using their respective in-house library preparation methods and NextGen sequencing platforms. Details can be found in Supporting Information file.

### Differential gene expression analysis and functional enrichment

DESeq2 v.1.34.0 (Love et al., 2014) was used to determine differentially expressed genes between turions and fronds. Shrunken log2-fold changes were generated using the DESeq2 shrinkage estimator using a Normal prior. To correct for multiple comparisons, p-values were adjusted using the Benjamini-Hochberg procedure. Expressed genes with a |log2-fold change| > 3 and an adjusted p-value < 0.05 were called as differentially expressed genes.

GOseq v.1.46.0 (Young *et al*., 2010) was used to perform functional enrichment of Gene Ontology (GO) terms for differentially expressed genes. GO.db v.3.14 was imported with GOseq to retrieve GO term annotations using the go-basic ontology release 2021-09-01. First, a probability weighting function was calculated for each gene to correct for gene length bias. Then, the wallenius approximation was used to calculate over- and under-expressed GO terms among differentially regulated genes. Finally, p-values were adjusted using the Benjamini-Hochberg procedure to correct for multiple comparisons.

### Quantitative analysis of triacylglycerol (TAG) and fatty acids (FAs)

Two clones of *Spirodela polyrhiza*, sp9509 and sp9512, were grown under conditions that favor either normal growth or the development of turions. Materials were harvested, freeze dried, and total lipids were then solvent extracted from the samples and separated by TLC using silica plates. The TAG-containing region of the TLC plates was isolated, and its constituent FAs were trans-esterified to corresponding Fatty Acid Methyl Esters (FAMEs). An internal standard of 17:0 FA was added to permit quantification of the FAME content, which was analyzed by GC-MS (Yu *et al*., 2014).

### Global mapping and comparison of cytosine methylation of genomic DNA in sp9512

High quality genomic DNA preparations from turion and frond tissues of sp9512 were prepared by bead beating of frozen plant samples and the Qiagen DNA isolation kit, according to the manufacturer’s protocol and reagents. DNA samples were sequenced using the ONT platform and cytosine methylation in the sequencing data were phase-called using the Megalodon pipeline and DMRs between turion and frond samples determined using Metilene (Jühling *et al*., 2016). Details of these methods and procedures can be found in Supporting Information file.

## Results

### Selection and genome assembly of a *S. polyrhiza* clone for turion studies

Leveraging a previously curated set of *S. polyrhiza* clones that were shown to exhibit a wide range of turion formation potentials (Kuehdorf *et al*., 2014), we identified a clone, sp9512, with rapid induction of turion formation upon decrease of the growth medium phosphate concentration to 2 μM (Fig. S1). Turions first became visible with sp9512 at 12 days after transfer to this inductive condition, while clone sp9509 requires about 22 days for the first appearance of turions. To support genomics studies with sp9512, we generated and characterized a high quality sp9512 genome assembly de novo that has similar or better metrics than the existing sp9509 reference genome (Figs. S3, S4),

### Improved protocol for turion RNA preparation enables high quality turion transcriptomics

Since extraction of high-quality nucleic acid from turions has posed a challenge for previous studies, we developed a new isolation technique that overcame this bottleneck. Three different homogenizing methods to process mature turion tissues before chemical extraction were compared: mortar and pestle, bead-beating, and a combination of mortar and beads (Fig. 1). We found a combination of tissue disruption by using mortar with bead-beating was necessary to improve RNA yield and quality from mature turions compared to when these procedures were used alone (Fig. S2B, C; Fig. S5). Total RNA was isolated from frond and turion samples of sp9512 and sequenced in replicates (see Supplemental Methods for details). Total number of raw reads obtained ranged from 20,559,476 to 48,056,644 with the constructed libraries. High-quality reads were mapped to either reference genomes with high alignment rates (Fig. S6, Table S1).

### Comparative analysis of global transcriptomes between mature turions and fronds

From the 18,403 protein-coding genes annotated in the sp9512 genome, 17,397 transcripts from the combined frond and turion transcriptome libraries were found, which represented 94.5% of its protein-encoding genes (Table S2). Of these, 14,137 protein-coding genes displayed less than a Log2-fold change of 2 between turion and frond transcripts (Fig. 2, Table S3). This result indicates that greater than 80% of the expressed *S. polyrhiza* genes showed less than a 4-fold difference in transcript abundance between these two developmental states. Thus, while turions are known to be dormant, most of their genes have transcript levels that are like in metabolically active frond tissues.

**Figure 2.**
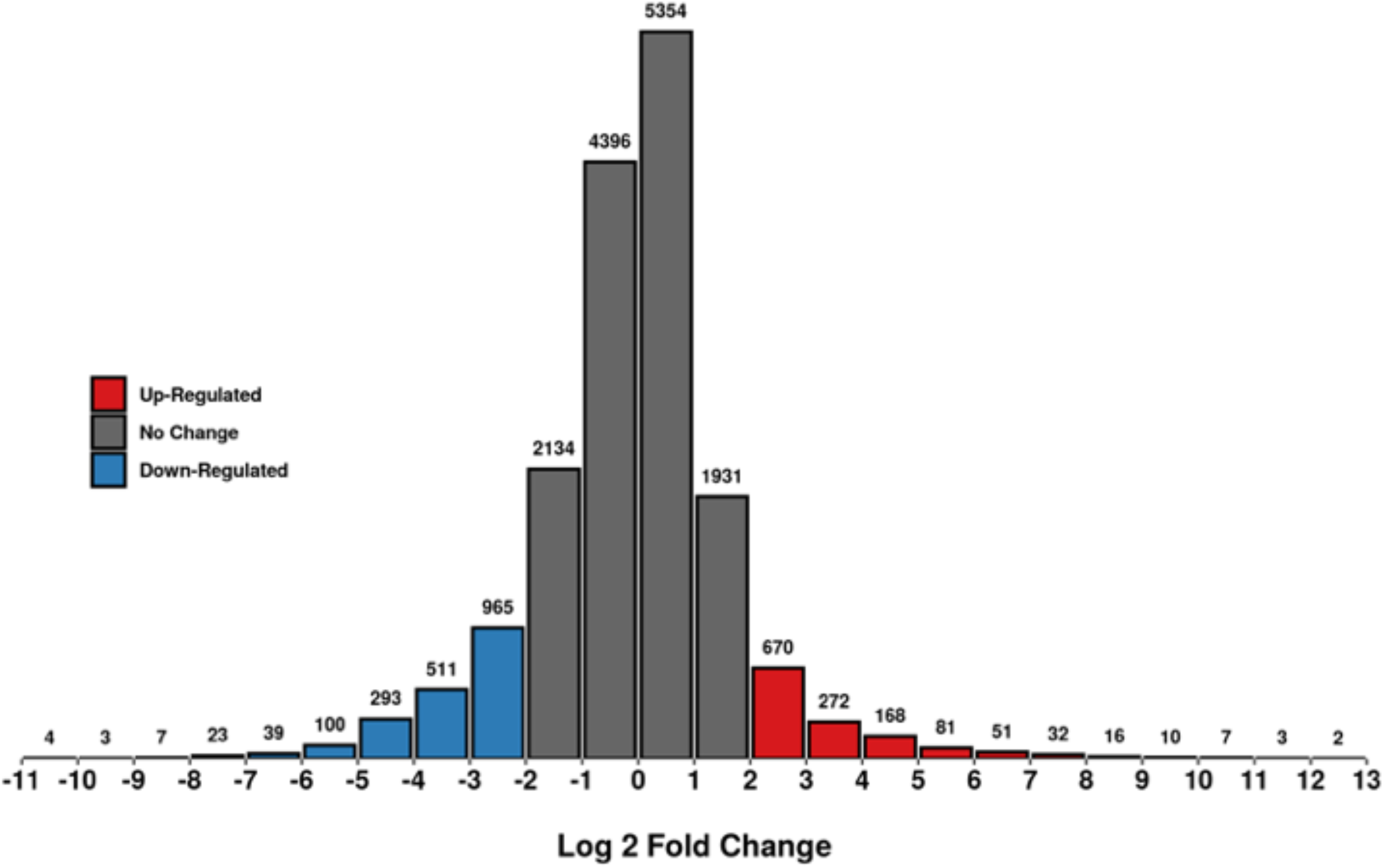
Histogram of Log2-Fold change in abundance of Sp9512 transcripts in turions vs. fronds. Differentially expressed genes are established at | log2-fold change | > 2. For GO analysis, a threshold of | log2-fold change | > 3 is used instead to increase the stringency for significant gene expression changes toward prediction of major pathways being altered. Red: significantly increased in turions; Blue: significantly decreased in turions; Black: not significant; Gray: significant but |log2-fold change| < 2; Significance = adjusted p-value < 0.05.

To reveal global changes in cellular pathways during turion formation, we carried out Gene Ontology (GO) functional enrichment analysis by curating genes with a |log2-fold change| > 3 (p. adjusted < 0.05) as a cutoff to increase the stringency for significance (Fig. 2). Through analysis of these genes by their GO terms, we found 4 major categories of pathways that are overrepresented by 8-fold or more in turions when compared to frond tissues (Fig. 3, Table S4). The top category with the largest number of affected genes and highest significance is related to stress responses, with many genes related to ABA signaling well-represented among this category (Table S5). Prominent among these are 2 genes encoding putative homologs to ABI5 (Abscisic acid-insensitive 5), a conserved leucine-zipper type transcription factor that mediates ABA-regulated gene expression (Li *et al*., 2022). In turions, the expression level of one of these ABI5-like genes is 39-fold higher, while the second one is 9-fold higher than in fronds. In addition, protein phosphatase 2C (PP2C)-encoding genes that could act as negative regulators of ABA signaling (Rodriguez *et al*., 1998) are also up-regulated up to 128-fold in turions. Strikingly, one of the most highly up-regulated transcription factors in turions (over 3,000-fold higher than in fronds) is a homolog to Arabidopsis MYB70, which belongs to the S22 subfamily in the large R2R3 type MYB transcription factor family (Persak & Pitzschke, 2014). Members of the S22 subfamily are substrates for phosphorylation by mitogen activated protein kinase 3 (MAPK3) and contain a trans-repressor domain containing the EAR (<u>E</u>thylene-responsive element binding factor-associated <u>A</u>mphiphilic <u>R</u>epression) motif for negative regulation of transcription. MYB70 could also interact with ABI5 to modulate ABA regulation of root development as well as seed germination in Arabidopsis (Wan *et al*., 2021). Furthermore, a MYC2-like turion transcription factor that can mediate gene expression changes in response to ABA, as well as crosstalk with other phytohormones such as Jasmonic acid (JA) and gibberellic acid (GA) in Arabidopsis (Hong *et al*., 2012), is expressed about 20-fold higher in turions.

**Figure 3.**
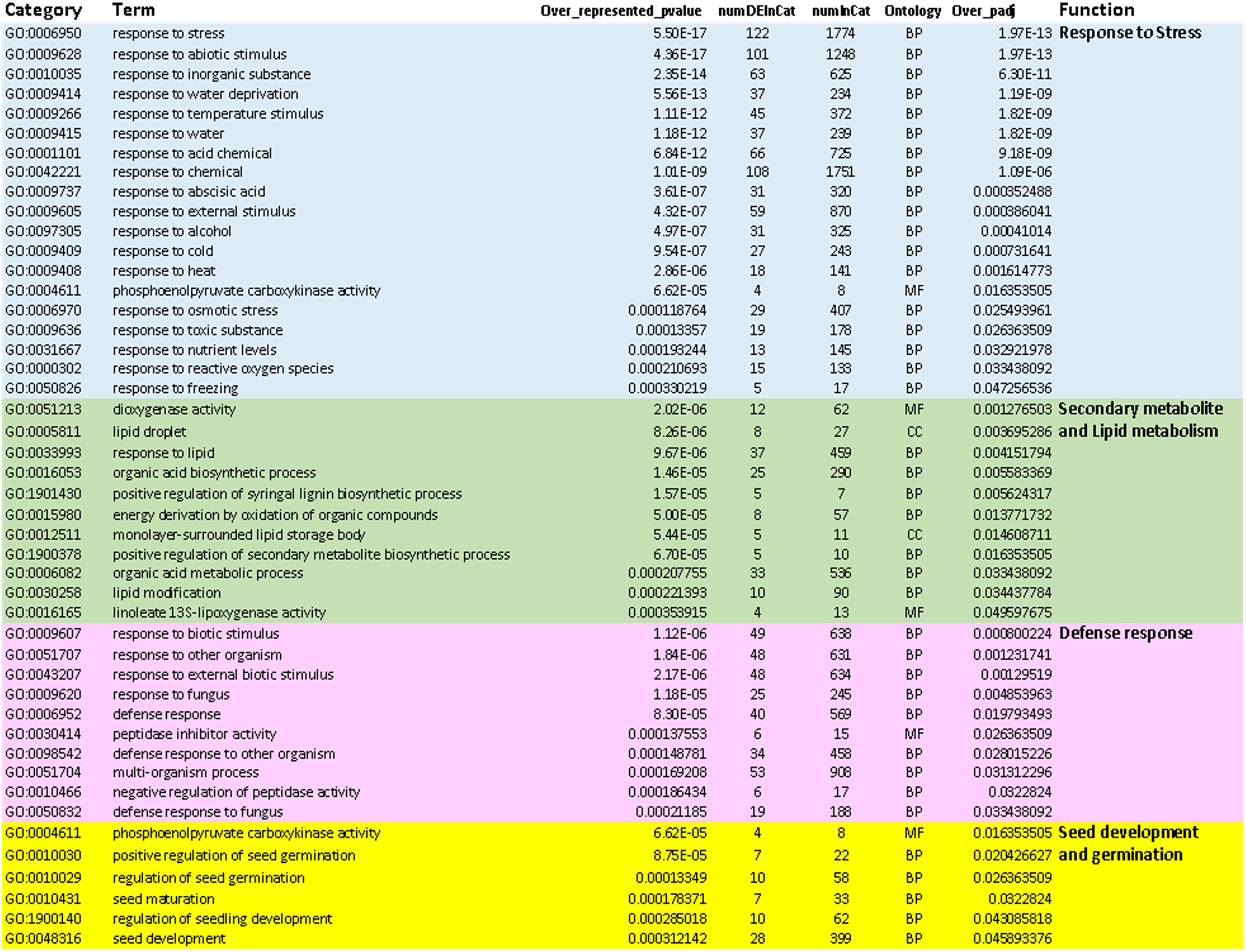
Overrepresented Gene Ontology (GO) terms for up-regulated genes in mature turions compared to frond tissues. A log2FC of 3 was used as a cutoff for inclusion of genes for this analysis.

Interestingly, a homolog to the Arabidopsis DOG1 (Delay of Germination-1) gene is induced over 100-fold in turions. DOG1 has recently been recognized as a master regulator of dormancy that works by enhancing ABA-dependency in part through its binding to PP2C and formation of a multiprotein complex with heme (Carilo-Barral *et al*., 2020). These findings provide support for the importance of ABA-mediated pathways in turion development and functions, as well as similarities to those related to seed dormancy and germination in land plants.

Other transcription factor types known to be involved in diverse stress responses are also heightened in their expression in turions. These include HSFs (heat-shock transcription factors), ICE1 (cold stress response), EREBPs (ethylene responsive element binding proteins) for tolerance to anoxia, and DREBPs (dehydration responsive element binding proteins) for drought stress response. Consistent with their increased expression, we found some of their target genes encoding HSP20 and HSP70, as well as a cold-regulated 413 (Cor413) membrane protein that could confer temperature stress protection (Zhou *et al*., 2018), being highly expressed in turions. Additional protection from DNA damage and endoplasmic reticulum dysfunction could be provided by up-regulation in turions of poly (ADP-ribose) polymerase (Aoyagi *et al*., 2021) and the conserved eukaryotic cell death repressor Bax Inhibitor-1 (BI-1) (Watanabe & Lam, 2009), respectively.

The second largest category of pathways that are significantly up regulated in turions is involved in secondary metabolite and phytohormone synthesis, as well as lipid biosynthesis and metabolism. As expected from the increase of anthocyanin in turions, a gene encoding 4-coumarate-CoA ligase (4CL5) that diverts secondary compounds from general phenylpropanoid metabolism to other branch pathways for secondary metabolites, is up-regulated by more than 30-fold. Consistent with the importance of ABA signaling for turion biogenesis, multiple enzymes for ABA biosynthesis are strongly increased at the transcript level. These include a gene encoding 9-cis-epoxycarotenoid dioxygenase that catalyzes the first step of ABA biosynthesis via cleavage of 9-*cis* xanthophylls to xanthoxin (Tan *et al*., 2003), and a homolog to Arabidopsis ABA2 that encodes a short-chain alcohol dehydrogenase which converts xanthoxin to abscisic aldehyde (Gonzalez-Guzman *et al*., 2002). Levels of these transcripts are increased by 80 to 100-fold in turions as compared to fronds. In addition to ABA biosynthesis, a gene encoding allene oxide cyclase, the key enzyme for JA biosynthesis (Zhang *et al*., 2002), is also increased by about 50-fold in turions. This indicates that JA responsive pathways may also play important roles in turion biology and is reminiscent of its involvement along with ABA in the regulation of seed germination (Zhang *et al*., 2002). Other classes of enzymes that showed significant increases in transcript levels are those involved in DNA and protein damage repair, as well as enzymes for cell wall modifications that could facilitate rapid growth during turion germination (Tables S3 to S5).

Included in this category of up-regulated metabolic pathways in turions are also two gene families involved in lipid accumulation and catabolic pathways that are of particular interest. Oleosins are important amphipathic proteins found at the surface of lipid droplets in seed tissues that protect triacylglycerides (TAGs) from TAG lipase attack (Graham, 2008). Overexpression of soybean oleosins in rice seeds can significantly increase oil content, indicating that oleosin levels can affect total oil content in plant tissues (Liu et al., 2013). We found 5 of the 7 oleosin-encoding genes in sp9512 are highly expressed in mature turions as compared to fronds (Tables 1, S5), along with two genes encoding proteins with homology to rubber elongation factor that may play a role in lipid droplet organization (Bröker *et al*., 2018). Sequence comparison of the sp9512 oleosin genes to other characterized plant oleosins indicate that 4 of the 5 highly expressed members in turions are more related to seed-specific oleosins in rice (Fig. S7; Table 1). Another group of lipid related genes which have heightened expression in turions are 3 different classes of lipases: GDSL lipase/acyl-hydrolases, lipolytic acyl-hydrolases (LAH), and a phospholipase A type lipase (Table 1; S5). Some of these could be involved in degradation of storage TAGs and fatty acids to support rapid frond development early in germination (Graham, 2008).

**Table 1.** *Spirodela polyrhiza* 9512 lipid metabolism genes and their relative expression in turions vs. fronds

| Pathway | Gene Number | Gene name | Turion/Frond (Log2Fold Change) |
| --- | --- | --- | --- |
| Synthesis | Sp9512.a02.1.g10270_p1 | SpWRI-a | -2.95 |
|  | Sp9512.a02.1.g10280_p1 | SpWRI-b | -3.04 |
|  | Sp9512.a02.13.g06050_p1 | SpWRI-c | -3.24 |
|  | Sp9512.a02.2.g08810_p1 | ACCase-a | -1.74 |
|  | Sp9512.a02.16.g04950_p1 | ACCase-b | -1.15 |
|  | Sp9512.a02.8.g02150_p1 | FUS3 | 0.31 |
|  | Sp9512.a02.6.g01920_p1 | FAX1 | 0.12 |
|  | Sp9512.a02.18.g02360_p1 | LEC2 | 0.71 |
|  | Sp9512.a02.19.g02960_p1 | TGD1 | -0.75 |
|  | Sp9512.a02.6.g07120_p1 | DGAT2-a | -1.75 |
|  | Sp9512.a02.10.g07200_p1 | DGAT2-b | -0.05 |
|  | Sp9512.a02.11.g02020_p1 | DGAT2-c | 1.13 |
|  | Sp9512.a02.7.g07790_p1 | PDGAT1 | 0.09 |
|  | Sp9512.a02.5.g08550_p1 | DGAT | 0.57 |
|  | Sp9512.a02.1.g06620_p1 | SDP1-a | -1.12 |
|  | Sp9512.a02.1.g06630_p1 | SDP1-b | -2.63 |
|  | Sp9512.a02.2.g12740_p1 | SDP1-c | 0.59 |
|  | Sp9512.a02.20.g03150_p1 | SEIPIN | -0.37 |
|  | Sp9512.a02.5.g03200_p1 | SpOLE1 | 4.60 |
|  | Sp9512.a02.5.g03230_p1 | SpOLE2 | 0 |
|  | Sp9512.a02.5.g08070_p1 | SpOLE3 | 5.82 |
|  | Sp9512.a02.11.g06030_p1 | SpOLE4 | 7.19 |
|  | Sp9512.a02.19.g04350_p1 | SpOLE5 | 6.61 |
|  | Sp9512.a02.20.g03650_p1 | SpOLE6 | 6.48 |
|  | Sp9512.a02.20.g04140_p1 | SpOLE7 | 2.76 |
| Degradation | Sp9512.a02.10.g05440_p1 | TAG Lipase | 0.81 |
|  | Sp9512.a02.2.g11710_p1 | SpLAH1 | 2.87 |
|  | Sp9512.a02.14.g00260_p1 | SpLAH2 | 0.25 |
|  | Sp9512.a02.14.g00290_p1 | SpLAH3 | 0.21 |
|  | Sp9512.a02.14.g00300_p1 | SpLAH4 | 4.79 |
|  | Sp9512.a02.16.g03490_p1 | SpLAH5 | -1.27 |
|  | Sp9512.a02.58.g00030_p1 | SpLAH6 | 3.38 |
|  | Sp9512.a02.1.g06260_p1 | Lipase (Class 3) | 0.81 |
|  | Sp9512.a02.1.g06280_p1 | Lipase (Class 3) | 1.04 |
|  | Sp9512.a02.2.g13030_p1 | Lipase 3 N-terminal | 0.78 |
|  | Sp9512.a02.3.g02770_p1 | Lipase (Class 3) | 0.29 |
|  | Sp9512.a02.2.g06210_p1 | Galactolipase DONGLE | 1.61 |
|  | Sp9512.a02.9.g04880_p1 | Phospholipase A1-Ibeta2 | 0.75 |
|  | Sp9512.a02.11.g03200_p1 | Phospholipase A1-Ibeta2 | 4.11 |
|  | Sp9512.a02.12.g02190_p1 | OB-Oil Body Lipase 1 | 3.24 |
|  | Sp9512.a02.2.g13280_p1 | GDSL-like Lipase | -0.97 |
|  | Sp9512.a02.2.g09270_p1 | GDSL-like Lipase | 5.21 |
|  | Sp9512.a02.3.g07720_p1 | GDSL esterase Lipase | 4.57 |
|  | Sp9512.a02.5.g05710_p1 | GDSL esterase Lipase | 2.96 |
|  | Sp9512.a02.5.g05700_p1 | GDSL esterase Lipase | 6.82 |

Relevant genes of interest involved in oil biosynthesis and degradation are curated from the assembled sp9512 genome and their transcript ration between turion vs. frond tissues shown as Log2Fold Change values. Values higher than 3 (eight-fold higher in turions) are highlighted in red and lower than -3 (eight-fold less) in teal.

As expected from the highly enriched starch content of mature turions (Fig. S2), we found the APL2 member of the conserved ADP-glucose pyrophosphorylase (AGPase) large subunit being highly expressed at more than 26-fold, with APL3 at 7-fold, higher than that of fronds (Table 2). In contrast, APL1 expression is more than 12-fold lower in turions, similar to the previous report with ABA treated fronds (Wang *et al*., 2014a). In addition to these significant changes, we found transcripts for a chloroplastic, α-glucan water dikinase that can facilitate branch-chain or crystalline starch degradation increased by more than 2,500-fold while a gene for β-amylase is more than 15-fold higher in turions as well (Table 2). Similar to the oleosins and lipases for lipid metabolism, these genes identified here present candidate genes that help to drive high levels of starch synthesis during turion maturation and their subsequent utilization upon germination.

**Table 2.** *Spirodela polyrhiza* 9512 starch metabolism genes and their relative expression in turions vs. fronds.

| Pathway | Gene Number | Gene name | Turion/Frond (Log2-Fold Change) |
| --- | --- | --- | --- |
| Synthesis | Sp9512.a02.1.g06970_p1 | AGPase Large subunit (SpAPL1) | -3.61 |
|  | Sp9512.a02.20.g05195_p1 | AGPase Large subunit (SpAPL2) | 4.74 |
|  | Sp9512.a02.6.g05320_p1 | AGPase Large subunit (SpAPL3) | 2.83 |
|  | Sp9512.a02.10.g06630_p1 | AGPase Small subunit | 0.56 |
|  | Sp9512.a02.3.g05980_p1 | Starch branching enzyme | -1.44 |
|  | Sp9512.a02.3.g06030_p1 | Granule bound starch synthase | 1.82 |
|  | Sp9512.a02.5.g04980_p1 | Starch synthase 1 | -1.14 |
|  | Sp9512.a02.12.g05830_p1 | Starch synthase 2 | 0.21 |
|  | Sp9512.a02.16.g06270_p1 | Starch synthase 3 | -1.66 |
|  | Sp9512.a02.16.g06300_p1 | Starch synthase 4 | -1.17 |
|  | Sp9512.a02.1.g06790_p1 | Isoamylase | -0.23 |
|  | Sp9512.a02.7.g08820_p1 | Isoamylase | 0.82 |
| Degradation | Sp9512.a02.5.g02610_p1 | Glucan water dikinase 2 (GWD2) | -1.06 |
|  | Sp9512.a02.6.g00030_p1 | Glucan water dikinase 3 (GWD3) | 11.31 |
|  | Sp9512.a02.12.g03390_p1 | Phosphoglucan water dikinase (PWD1) | -0.51 |
|  | Sp9512.a02.18.g00670_p1 | Alpha glucosidase | -0.75 |
|  | Sp9512.a02.15.g04080_p1 | Alpha amylase | -0.48 |
|  | Sp9512.a02.1.g07640_p1 | Beta amylase | 0.97 |
|  | Sp9512.a02.8.g06570_p1 | Beta amylase | 0.50 |
|  | Sp9512.a02.14.g01100_p1 | Beta amylase | 3.94 |

Relevant genes of interest involved in starch biosynthesis and degradation are curated from the assembled sp9512 genome and their transcript ration between turion vs. frond tissues shown as Log2-Fold Change values. Values higher than 3 (eight-fold higher in turions) are highlighted in red and lower than -3 (eight-fold less) in teal.

The third category of GO terms that are significantly enriched in turions is defense response to biotic agents. While many of the genes involved are shared with those in the first category of Stress Responses (Tables S3, S4), we note heightened expression of several receptor-like serine and threonine protein kinases that could mediate defense to bacterial pathogens. In addition, two genes that could mediate defense to oomycetes, one encoding a multi-antimicrobial extrusion (MATE) family member while another is related to the Arabidopsis LURP1 protein that can mediate salicylic acid activation of defense to the oomycete *Hyaloperonospora parasitica* (Dobritzsch *et al*., 2016; Knoth & Eulgem, 2008), are upregulated by about 10-fold in turions. Also, transcripts for a gene encoding an endochitinase that can degrade fungal cell walls, and for a receptor kinase with a lysM (Lysine Motif) domain that likely mediates cellular response to chitin, are highly increased in turions. Likewise, 6 genes with homologies to the MiAMP1 class of antimicrobial peptides that has recently been found to be upregulated in duckweeds upon bacterial pathogen challenge (Baggs *et al*., 2022) are increased in expression by 10- to 47-fold. Finally, we note that 4 genes in the phytocystatin family among its 27 members curated in the sp9512 genome are highly expressed in turions. Their increase varies from 128- to 8-fold higher and indicate that these potent cysteine protease inhibitors could play important roles for protection from insect herbivores or other pathogens, such as fungi, which may utilize cysteine proteases to digest or colonize turion tissues. Together, these results indicate turions have hyperactivated surveillance to guard against various types of potential invaders.

Interestingly, the fourth category of turion up-regulated genes are related to seed development and germination. Here, regulators of the response to ABA in seed maturation and embryogenesis (Ali *et al*., 2021) are well-represented (Table S5). In particular, transcripts for a homolog to Mother of FT and TFL1 (MFT), a phosphatidylethanolamine-binding protein that directly regulates ABI3 and ABI5 transcription factors (Xi *et al*., 2010), is increased by 13-fold in turion vs. frond tissues. Through integrating the ABA and GA signaling pathways, MFT mediates seed germination control in model plants such as Arabidopsis. In addition to increases of ABI5 expression noted earlier, two additional transcription factors involved in seed-related development are also upregulated in turions: one with homology to AtMYB5 that regulates mucilage biosynthesis in the developing seed coat (Li *et al*., 2009) and a MADS class transcription factor related to those involved in endosperm development such as AGL62 (Kang *et al*., 2008), are 9-fold and 11-fold higher in turions, respectively. Finally, one of the six genes encoding an A-subunit of the trimeric Nuclear Factor (NF)-Y transcription factor in sp9512 is upregulated more than 20-fold in turions. Members of this NF-Y subunit have been shown to play crucial roles in early embryogenesis as well as mediate ABA sensitivity in the vegetative-to-embryonic transition (Zhao *et al*., 2017). While genesis of the metabolically dormant turion has been thought to be more like vegetative bud formation based on morphological considerations (Adamec, 2018), these molecular data indicate a striking similarity at the transcription factor level to that mobilized for seed formation and maturation of the zygotic embryo during sexual reproduction. This analogy extends to the activation of similar types of LEA (Late Embryogenesis Abundant) genes upon turion induction, which has been well studied as widely conserved proteins for stress tolerance from algae to land plants and as cellular protectants during seed formation (Hundertmark & Hincha, 2018). LEA proteins are relatively small (10-30 kDa), hydrophilic, and unstructured proteins that appear to function as “buffers” to prevent proteins from aggregating in the cell, which often happens upon abiotic and biotic stresses. Five types of LEA proteins are defined by distinct repeated sequence motifs and together with Dehydrins and Seed Maturation Proteins (SMPs) made up the 7 types of LEA genes that we could curate from the *S. polyrhiza* genome (Fig. S8, Table S5). Among the 50 LEA genes that we identified in the sp9512 genome, 13 are upregulated in turions and varied from 11- to over 2,000-fold. Interestingly, 5 out of the top 6 highly expressed LEA genes in turions are most homologous in sequence to seed-specific LEAs in rye (Ding *et al*., 2021), suggesting that similar preference for those types of LEAs with distinct functions is shared between seeds and turions (Fig. S8).

Finally, we note heightened expression for 4 genes encoding the enzyme ATP-dependent PEPCK (phosphoenolpyruvate carboxykinase), which ranged from 15- to over 100-fold higher in turions. This enzyme is known to play an important role during seed germination by channeling high energy intermediates from fatty acid degradation to provide sugars through gluconeogenesis for germinating seedlings (Graham, 2008; Rojas *et al*., 2019). In this regard, it is interesting to note that genes encoding malate dehydrogenase and a malic enzyme with decarboxylating (NADP+) activity also have increased expression in turions. These enzymes play important roles in TAG degradation via the glyoxysome during early seedling development in Arabidopsis and could similarly work together with PEPCK to provide much needed energy during turion germination (Graham, 2008; Yokochi *et al*., 2020).

### Major gene ontology terms that are under-represented in turions compared to that in fronds

Since turions are known to be in a quiescent state until after the overwintering process to break dormancy, we anticipate transcripts and processes that normally require a lot of energy expenditure to maintain but are dispensable during early germination would be under-represented in turions as compared to fronds. We examined the GO terms for turion transcripts that decreased by 8-fold or more when compared to that in fronds (Log2FC ≥ 3 in fronds vs turions; Table S6) and found the major categories are related to cell division, DNA replication, and cytoskeleton related processes (Fig. S9). Similarly, some genes that participate in organelle fission as well as for developmental transition in either dark (skotomorphogenesis) or light (photomorphogenesis) are decreased. The next group of genes that are significantly down regulated in turions are those involved in ion transport, notably those for anions and nitrate. Finally, genes involved in cell wall modifications such as suberin production are also repressed.

Instead of globally screening for down-regulated pathways, we also examined two nuclear gene families that encode highly expressed proteins in plants: the small subunit of RubisCo enzyme (RbcS), the most abundant enzyme on earth, and the light-harvesting antenna (LHC) proteins. We curated 7 RbcS and 16 LHC encoding genes in the genome of sp9512 and found that 4 of the 7 RbcS genes are among the top seven transcripts most strongly down-regulated in turions in comparison to fronds, with the top one at more than 1,700-fold lower expression (Table S7). None of the members in these two gene families showed higher expression in turions, while almost all members displayed a decrease of more than 4-fold. Thus, it appeared that much lower levels of transcripts for these two gene families are being maintained in the turion since they are likely not needed during the dormant stage with translation activity essentially arrested.

### PCR-based Validation of DEGs and comparative analysis of transcript induction rates between clones

To validate some of the predicted DEGs between turion and frond tissues, we compared transcript quantification by reverse transcriptase qRT-PCR and DESeq2 normalized read counts from the transcriptome datasets for three genes of interest that span different ranges of read counts (Fig. S10, Table S9). We also tested 9 additional turion up-regulated genes using end-point RT-PCR assay, while including one RbcS gene as a turion-repressed transcript and one actin gene as a constitutive control. Our results uniformly showed clear transcript detection for the 9 target genes in turion but not frond samples, while the RbcS gene and actin control displayed expected amplification patterns (Fig. S11). These results validate the quality of our transcriptome datasets and support our conclusions on global transcriptome differences between turions and fronds.

Based on the significant difference in apparent turion induction rates between clones sp9512 (12 days) and sp9509 (22 days), we expect many of the turion-specific genes may exhibit markedly different rate of activation in these clones upon low phosphate trigger. Profiling transcriptomes from these two clones at various points between 7 and 25 days post-transfer to low phosphate medium, we found this is indeed the case for many of the turion-specific genes such as those encoding MYB70, PP2C-1, GWD3 and OFT (Fig. S12), In contrast, several other turion-up genes such as OCT4, PARP3 and BI-1 showed similar induction rates between the two clones, suggesting they are induced in the frond tissues as well under low phosphate.

### Characterization of triacylglyceride content and fatty acid composition differences in turions and fronds

One interesting observation from our transcriptomics study is the discovery of oleosin genes being highly expressed in turions (Table 1). This observation is unexpected since previous microscopy work with early turion biogenesis suggested that lipid droplet numbers do not change significantly in abundance from those in frond tissues (Smart & Trewavas, 1983b). We thus carried out quantitative analysis of the TAG content in turions and fronds (Fig. 4A) and determined the composition of TAG-associated fatty acids (FAs) via gas chromatography-mass spectrometry (GC-MS) (Fig. 4B). For our analysis, we included frond and turion samples from both clones sp9509 and sp9512 to evaluate any potential intraspecific variation. Results from this study show that turions consistently showed between 2.6- and 2.9-fold increases in TAG as compared to fronds for both clones (Tukey’s multiple comparisons test, P<0.05). Staining of the thin layer chromatography (TLC) plates with primuline revealed a higher level of free FA (FFA) in the lanes containing lipids from fronds, when compared to lanes containing turion lipids (Fig. S13). The accumulation of FFA suggests that the rate of fatty acid synthesis may outpace cellular demand in rapidly growing fronds. We therefore quantified the level of FFA in each sample. The data show that fronds of sp9509 and sp9512 contain between 3.8 and 4.2-fold higher FFA (Tukey’s multiple comparisons test, P<0.05) compared to turions from the same clones (Fig. 4C). Compositional FA analysis of the TAG pool showed that fronds of sp9509 and sp9512 have significantly higher levels of 16:0, 18:0, 22:0 and 26:0 saturated FA than corresponding turions from the same clones (Fig. 4B), while the turions had more than four times the level of the di-unsaturated linoleic acid. In contrast, the FA profiles from the FFA pool of fronds and turions showed no significant difference in their FA compositions, except for small differences in 16:0 which was significantly higher in fronds than turions and 24:0 which was significantly higher in turions than in fronds (Fig. 4D). These changes prompted us to analyze the total FA (TFA) profile of fronds and turions. We observed that turions show a 50% or greater decrease of the level of TFA relative to growing fronds (Fig. 4E). The data also show significant (Tukey’s multiple comparisons test, P<0.05) changes in the distribution of polyunsaturated fatty acids between fronds and turions (Fig. 4F). In Fronds, the predominant polyunsaturated fatty acid is 18:3, whereas in turions, 18:2 predominates. In sum, mature turions contain greater than a 2.5-fold increase in TAG levels and a corresponding decrease in the TFA and FFA levels relative to fronds of the same clones, along with a reduction of 18:3 and increase in 18:2. Turions also show a large increase in the level of the di-unsaturated FA linoleic acid and a corresponding decrease in the levels of saturated 16:0, 18:0, 22:0 and 26:0 FA in its TAG relative to those from fronds.

**Figure 4.**
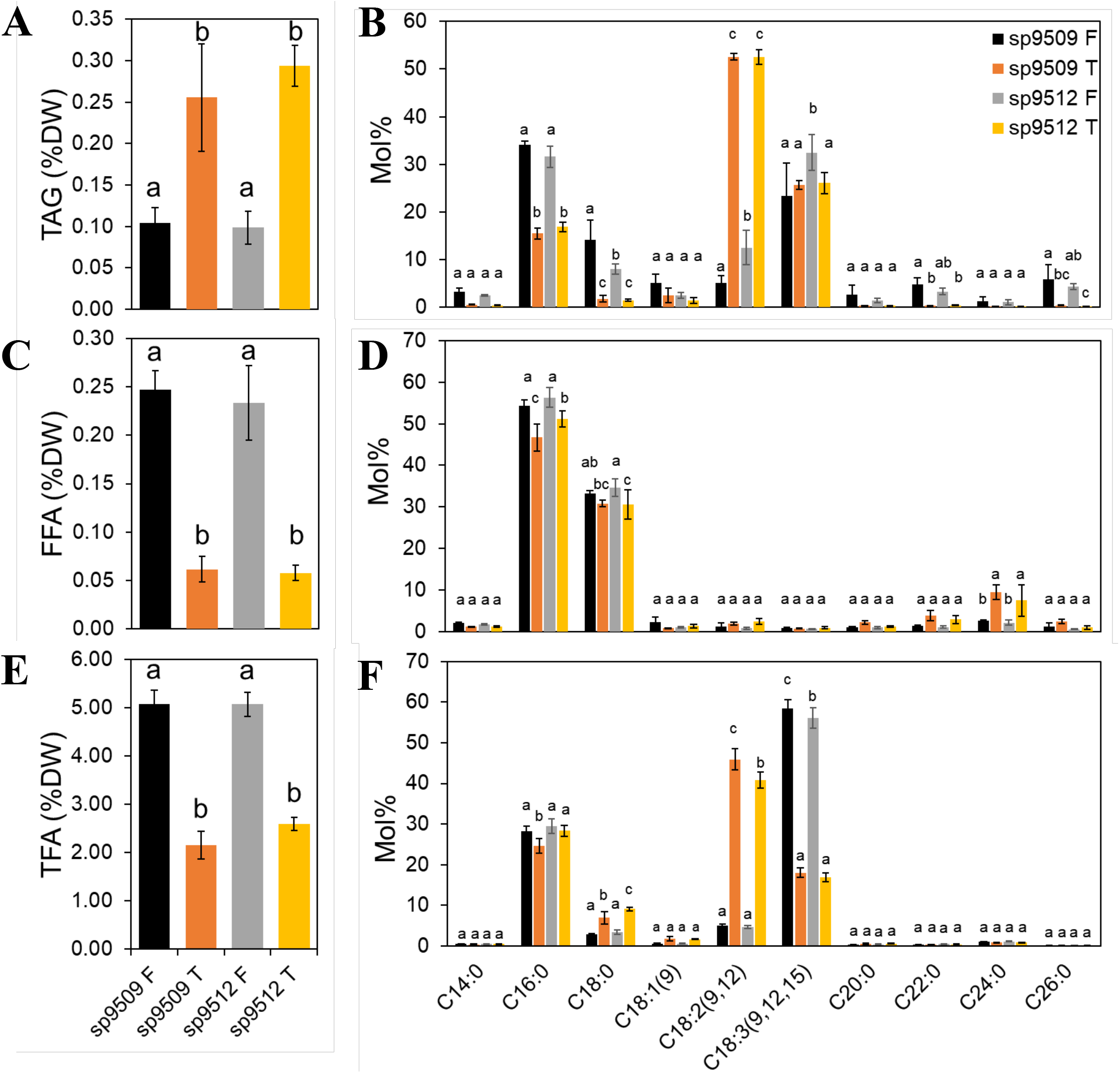
Analysis of TAG, FFA and TFA from *Spirodela polyrhiza*. (A) TAG content of fronds and turion of *S. polyrhiza* (sp9509 and sp9512). (B) Fatty acids profile of TAG in fronds and turions of *S. polyrhiza* (sp9509 and sp9512). (C) FFA content of fronds and turion of *S. polyrhiza* (sp9509 and sp9512). (D) Fatty acid profile of FFA in fronds and turions of *S. polyrhiza* (sp9509 and sp9512). (E) TFA content of fronds and turion of *S. polyrhiza* (sp9509 and sp9512). (F) Profile of TFA in fronds and turions of *S. polyrhiza* (sp9509 and sp9512). F: Frond, T: Turion.

### Global changes in cytosine methylation patterns upon turion formation mirrors similar genomic alterations observed during seed maturation

Epigenetic regulation via reversible cytosine methylation is one mechanism involved in facilitating developmental transition in both animals and plants (Law & Jacobsen, 2010). We wanted to examine whether the frond to turion developmental transition could involve changes in DNA methylation status in the genome as well. To examine the global status of cytosine methylation, whole genome 5-methylcytosine (5mC) mapping and quantification in frond and turion genomic DNA was carried out using the Megalodon pipeline with ONT (Liu *et al*., 2021). As we have previously reported using bisulfite-seq (Michael et al., 2017), *S. polyrhiza* fronds have very low overall 5mC content (<10%) compared to other model species such as Arabidopsis (≥30%) (Fig. 5). Separating the three types of target site context (CpG, CHG, and CHH), and examining different regions that contain either genic or transposable elements (TEs, Fig. 5D), we found the transition to turion correlated with global hypermethylation of TE-localized CHG and CHH sites, although significant increases of these methylation target sites can also be seen in the genic regions (Fig. 5A and 5B). Globally, a slight decrease in methylation is only observed in CpG sites inside genic regions (Fig. 5A). This is reflected in scanning through each of the 20 assembled sp9512 chromosomes, using a 100-Kb bin, with an example shown for chr. 17 (Fig. 5C). This chromosome is the only one out of the 20 that showed two peaks of hypomethylation in CpG sites on turion DNA vs. frond DNA. Both CHG and CHH sites only displayed increases in turion DNA when compared to the corresponding sites in fronds. While we did not find a strict correlation at a global level between differentially expressed genes and predicted hypomethylation or hypermethylation, we did uncover several instances where corresponding CpG methylation level changes were observed at promoter regions of genes that were either activated or repressed in turion tissues relative to those of fronds (some examples shown in Fig. S14). In addition, the global increase in TE methylation at the CHH sites found in turion DNA is similar to that in studies involving global methylation changes during seed maturation and dormancy onset in Arabidopsis (Lin *et al*., 2017; Kawakatsu *et al*., 2017). As discussed previously (Lin *et al*., 2017, Kawakatsu *et al*., 2017), this hypermethylation of TEs in the genome likely helps to ensure that they remain inactive during dormancy. In addition, the increase in methylation overall in the genome could help guard against DNA damage that the dormant turion may be more susceptible to when its enzymes for DNA repair are less active.

**Figure 5.**
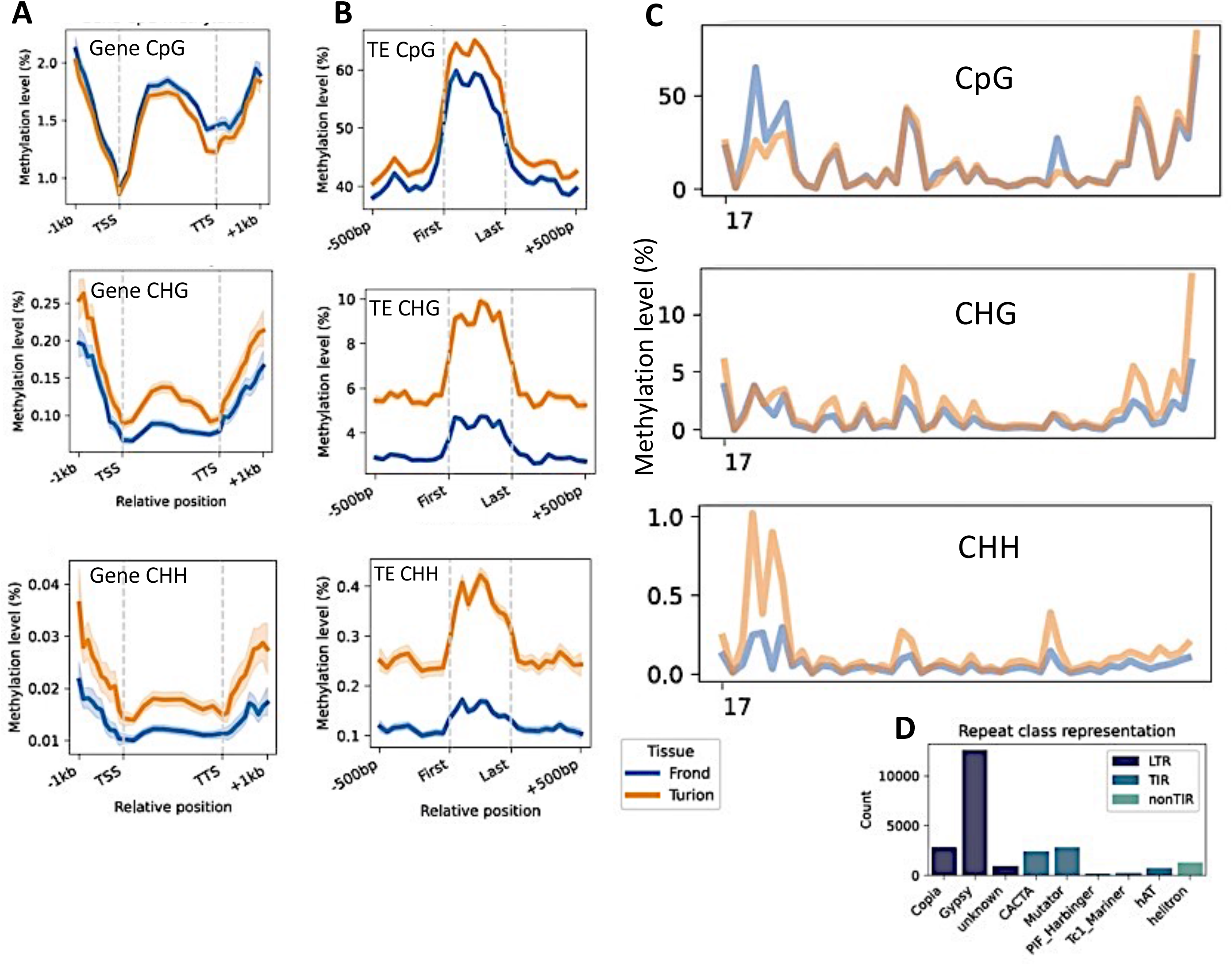
Genomic cytosine methylation levels in frond and turion DNA. A) Average methylation levels across gene bodies, 99% confidence intervals shown. B) Average methylation levels across transposable elements (TEs). C) Average methylation levels of 100-Kb bins across Chr. 17. D) Repeat classes and their abundance in the sp9512 genome. CpG, CHG and CHH residues are separately displayed as shown.

## Discussion

### Comparative transcriptomics between frond and turion tissues reveals genes and pathways that contribute to duckweed’s survival strategies

As small, floating macrophytes on water bodies, duckweeds’ major competitors for growth and survival are algae that thrive in the water column. Under warm temperatures and with sufficient nutrients, duckweed can rapidly cover the water surface through its fast growth and ready access to sunlight. This results in effective shading of the algae below and can provide a competitive edge for duckweed. As the season progresses to autumn, however, nutrients can become depleted while temperature also decreases in temperate regions. These environmental conditions often combine to trigger, in species such as *S. polyrhiza*, the formation of turions as propagules that settle to the bottom of the water bodies where they remain until better growth conditions return (Figure 6). Thus, turions serve the function as “escape pods” that ferry a clonal derivative of the duckweed frond cluster to relative safety where it can lie in a dormant state. After it has overwintered there through the cold months of winter, turion germination can be triggered by the red-light absorbing chromophore phytochrome (Appenroth et al., 2001) with the re-emergence of the frond and root, concomitant with its increase in buoyancy as it resurfaces (Figure 6). During this time, the ability to quickly resume active photosynthetic growth capacity is important to drive the rapid growth of duckweed again. Indeed, measurement of dark respiration and photosystem II (PSII) fluorescence quenching in overwintering and germinating *S. polyrhiza* turions showed transition from low photochemical efficiency, with the majority of the PSII reaction centers uncoupled from the electron transfer chain to full recovery by two days after germination (Oláh et al., 2017). While some of the biochemical events during the overwintering and germination processes of turions have been described, such as the activation of starch metabolism (Appenroth & Ziegler, 2008), few details of the global response are known. Our present study provides the first comprehensive description of the genes and pathways that are strongly affected during the transition from frond to turion in *S. polyrhiza*. The results not only delineate a set of turion-specific genes that are activated under low phosphate concentrations, but they also revealed that turions store a large variety of mRNA transcripts in addition to its abundant starch. With these stored transcripts, the germinating turion in the Spring should be able to quickly reactivate essential pathways by the resumption of translation activities. The abundant starch that is stored in turions could fuel growth in the early stages of germination (Fig. S15; Table 2), while the TAG in lipid droplets described here can provide an additional source of reduced carbon (Fig. S16) to sustain growth and development of fronds before they become fully active and photosynthetically competent. The reduced content of TFA and FFA in turions along with large decrease in the levels of 18:3 in TFA is consistent with the loss of photosynthetic membranes during the transition from growth as fronds to more dormant tissue such as turions, while the increased levels of unsaturated fatty acids in turion TAG is likely advantageous for their incorporation into new membranes that have sufficient membrane fluidity to maintain function in cold, post-germinative growth conditions. Our work thus helps to lay the foundation for molecular dissection of the genes that may be required for turion development, such as the highly turion-specific MYB70 and PP2C-1 genes (Fig. S12), as well as identified specific gene targets to manipulate starch and oil biosynthesis (Tables 1 and 2) in *S. polyrhiza* and engineer optimized sustainable feedstocks to augment traditional crops.

**Figure 6.**
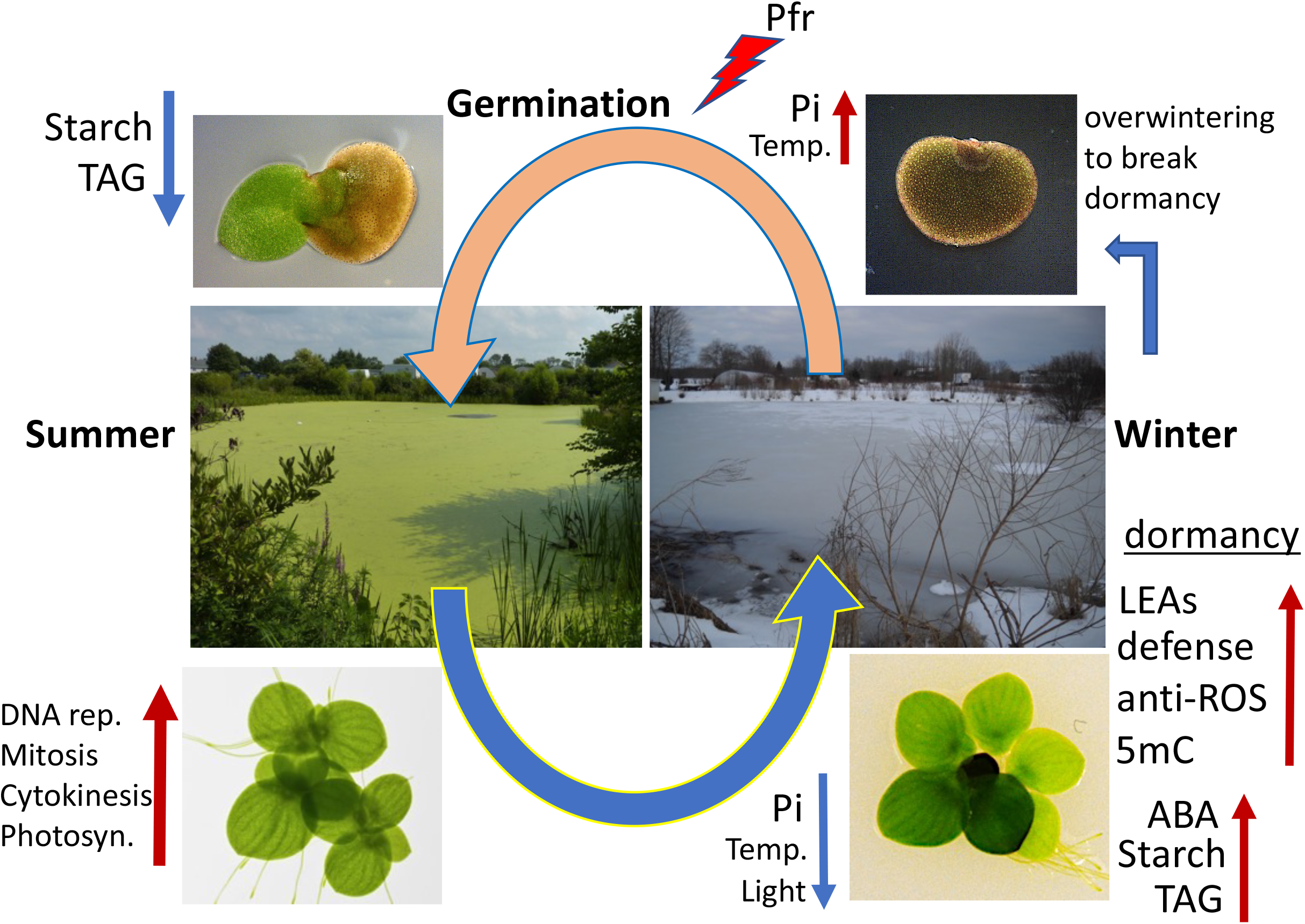
Turions as “escape pods” for *Spirodela polyrhiza* to survive environmental stresses. Current model of the apparently strategy that has evolved in the Greater Duckweed to trigger developmental transition at the meristematic tissue to produce turion instead of frond. The conditions in autumn (decrease in phosphate, temperature and daylight) trigger formation of turions, which became dormant as they mature and accumulate starch and TAG. As turions overwinter at the bottom of the water body, dormancy breaks toward Spring and the far-red absorbing (active) form of phytochrome (Pfr) triggers the turion to begin germination.Metabolizing the stored starch and TAG in the germinating turion then fuel the growth of the young fronds and roots that are emerging from the turion so that the young fronds can grow rapidly while re-establishing all the proteins needed for efficient photosynthesis to provide the energy needed to rebuild an actively growing frond cluster as quickly as possible.

### Similarities of turion-specific genes/pathways to seed development in plants and hibernation in animals

Finally, our results indicate pathways and genes that are utilized for zygotic seed development may have been re-tooled for use in asexual production of turions in the Greater Duckweed. The strategy of transcript storage revealed by our turion transcriptome profiling study also has interesting corollary to findings from animal hibernation studies. Like dormancy in plants, mammalian hibernation involves drastic reductions in normal metabolic activity to 5% or less of that in the resting state (Morin & Storey, 2009). A recent transcriptomics study with three different tissues during the hibernation cycle in grizzly bears dissects the specific changes in transcript abundance between the adipose, liver, and skeletal muscle tissues (Jansen *et al*., 2019) during key points in the hibernation /active/hyperphagia cycle. Among these tissues, 80% or more of the transcripts expressed under the active stage are not significantly altered in the hibernation stage, with the adipose tissue showing the most differentially expressed genes. Of these, GO enrichment analysis also revealed the metabolic pathways for gluconeogenesis and lipid metabolism are the top categories being rewired during transition to hibernation in animals as well. While the specific identity of the enzymes and genes that are affected often differ between animal and plant systems, due in a large part to the divergent specificities of the physiology and biochemistry that are involved across kingdoms, the strategy is remarkably similar in terms of the increased synthesis and storage of polyunsaturated fatty acids before and during dormancy or torpor. Future studies of the details in genetic and potential epigenetic pathways, such as histone modifications and long non-coding RNAs (Morin & Storey, 2009; Jansen *et al*., 2019), in both animal and plant dormancy systems would help to reveal the molecular pathways for coordinated gene expression control underlying this conserved strategy for survival in biology.

## Acknowledgements

Duckweed research at the Lam and Shanklin laboratories is supported in part by a grant from the Department of Energy (DE-SC0018244). The Lam lab is also supported by a Hatch project (#12116), and a Multi-State Capacity project (#NJ12710) from the New Jersey Agricultural Experiment Station at Rutgers University. J.S. was supported in part by the US DOE Office of Basic Energy Sciences (DE-SC0012704). This work in the Michael lab was supported by the Tang Genomics Fund.

## Author contributions

B.P. performed all the experiments for turion induction, physiological studies, and nucleic acid isolation for transcriptomics studies; B.A., K.C., N.H. carried out HMW genomic DNA isolation from tissues, DNA library construction, and Nanopore sequencing and genome assembly as well as transcript mapping analyses; N.H., B.P., and K.A. carried out the bioinformatics analyses for differential gene expression; T.P.M. carried out detailed genome comparisons between the three *S. polyrhiza* reference genomes; A.A. and B.A. did the 5-methylcytosine mapping and differential analysis of transcripts and methylation; Y.L. and J. S. performed the lipid analyses of plant tissues; B.P. and E.L. designed the experimental setup; E.L., B.P., K.A., T.P.M., A.A. and J.S. wrote the manuscript draft; all authors provided editing and final approval for submission.

## Data and Code Availability

Genomic and RNA-seq reads related to the sp9512 frond and turion study can be found in NCBI under BioProject: PRJNA858442 and Biosample: SAMN29716464

## Supporting information

Supplemental information and tables attached

