## Supplemental File and Figures for "Genomics of turions from the Greater Duckweed reveal its pathways for dormancy and reemergence strategy"

**Pertinent Information of Turion Biology in *Spirodela polyrhiza***

While sexual reproduction in plants is usually linked to seasonal cues to initiate the production of seeds with dormant embryos as propagules for the subsequent generation, aquatic plants that prefer clonal propagation for normal growth adapts to these environmental cues have evolved the strategy of turion production for this purpose. After their formation from the meristem pocket, turions emerge from the mother frond cluster (Fig. 1A) and sink to the bottom of water bodies in autumn after maturation and detachment, where they wait for the proper environmental conditions for germination and reemergence to the surface. To characterize the metabolic and structural changes of the induced fronds in *S. polyrhiza*, the phytohormone abscisic acid (ABA) is a useful tool as a convenient chemical trigger for turion production, and dose dependence of ABA concentration as well as the time course for this induction process have been defined (Smart & Trewavas, 1983a). Using radioisotope-labeled precursors, DNA synthesis was shown to be inhibited within several hours after ABA addition, followed by shut down of protein synthesis within one day after treatment (Smart & Trewavas, 1984). In contrast, RNA synthesis did not cease until 3 days after ABA addition, which coincided with the time that the developing turion first became visible under these conditions. Whereas young vegetative fronds develop prominent mesophyll and vascular tissues, cells of the turion-forming, embryo-like cells accumulate starch and do not show any obvious aerenchyma. These embryo-like structures further develop into mature turions with most cells accumulating high levels of plastidic starch and tannins. Together with the decreased water content in turions of about 70-80%, in contrast to the 90-95% water content for fronds (Landolt, 1986), these physiological characteristics likely contribute to the lower buoyancy as well as dormancy of mature turions.

**Sequencing and assembly of the sp9512 genome.** To facilitate the study of turion biology, the identification of an accession of *S. polyrhiza* with rapid induction kinetics as well as high turion yield was desirable. By comparing five different clones selected from a set of 36 previously shown to display a wide range of STY (Kuehdorf *et al*., 2014), we found the turion formation rate, as measured by the first day of visible turions upon phosphate limitation in the growth media, varied from 12 (clone sp9512) to 22 (clone sp9509) days (Fig. S1). These differences largely correlated with their reported variations in STY, with clones of higher STY displaying a shorter initiation time following the low nutrient trigger. Comparison of physical and biochemical parameters between fronds and turions from sp9512 and sp9509 showed that turions have lower chlorophyll content and higher starch (Fig. S2), consistent with previous reports (Landolt, 1986). For our study, we thus chose the sp9512 clone as our standard strain and used transfer to low phosphate (2 µM) media as the physiological turion induction trigger.

We generated a high-quality reference genome for sp9512 to complement the published reference genome for sp9509 (Hoang *et al*., 2018) and to aid molecular studies of turion formation in this genetic background. Clone sp9512 was sequenced with Oxford Nanopore Technologies (ONT) long reads and assembled into 200 contigs with N50 length of 4.5 megabases (Mb) (Fig. S3). The sp9512 contig assembly (sp9512v1) was then scaffolded based on the chromosome-resolved sp9509 reference (Hoang *et al*., 2018) resulting in a final genome assembly (sp9512v2) with a total length of 140 Mb and N50 scaffold length of 7.8 Mb (Fig. S3). Gene prediction resulted in 18,402 protein coding genes, which is similar to that found in both high-quality genomes of sp9509 and sp7498 (Hoang *et al*., 2018; Harkess *et al*., 2021). In addition, 14.5% (2,751) of the genes were found in tandem duplications (TD) representing 829 families, while 54% of genes were retained as multiple paralogs in syntenic blocks as a result of likely past whole genome duplication (WGD) events (Figs. S3, S4). As expected from the known high degree of intraspecific sequence conservation in *S. polyrhiza* (Michael et al.,

2017; Xu et al., 2019), dotplot analysis between sp9512v2 and the other two published reference genomes from sp9509 and sp7498 clones revealed high degrees of synteny (Fig. S4). Two recent WGDs have been identified in *S. polyrhiza* (4:4 syntenic pattern), and it has been suggested that *S. polyrhiza* does not have the tau WGD shared across other monocots (Wang et al., 2014b). However, recent evidence from additional high-quality duckweed genomes for Lemna and Wolffia, which do not have the more recent WGD events, suggest that the tau WGD might be obscured by the most recent WGD in *S. polyrhiza* (Abramson et al., 2021). To test if *S. polyrhiza* may have experienced more fractionation after the recent WGD, we modified the analysis of syntenic blocks to account for larger more fractionated blocks and found evidence for the tau WGD in sp9512 (Fig. S4A, D). Consistent with this finding, despite that greater than 99% of genes are found in syntenic blocks between sp9512 and sp9509 (or sp7498), 8-9% of genes have been differentially fractionated after the two recent WGDs (Fig. S4). Thus, even though *S. polyrhiza* has a low level of nucleotide variations between clones, the genome structure is significantly variable across the three reference quality assemblies analyzed here, which could play a role in the differential turion formation characteristics observed between the clones (Kuehdorf *et al., 2014)*.

**Comparative transcriptomic analysis between clones with divergent STY provides insights for specificity of turion associated genes.** To further characterize the specificity for some of the most highly expressed genes in turions, we carried out time course studies over 25 days after shifting fronds of sp9512 and sp9509 to media with low phosphate. Since we found that these two clones differ in their turion initiation rate by more than 1 week, we hypothesize that activation of genes which are tightly associated with turion biogenesis would also be delayed in the sp9509 clone upon low phosphate treatment. Transcriptome libraries were generated with tissues of these two clones of *S. polyrhiza* at four time points between 7 and 25 days after shifting into media with low phosphate for turion induction. With sp9512, turions first became visible at ~12 days while for sp9509, ~22 days are needed (Fig. S1). cDNA sequence reads from each of these clones were mapped to the sp9512 chromosome-resolved genome assembly and transcripts were sorted to curate genes with low expression (normalized read counts from 0 to 30) at 7 days post transfer to low phosphate media but with high expression at 25 days post transfer. Using this approach, we focused our comparison of induction kinetics on genes that have low background expression in fronds to minimize the complex mixture of tissue types in our samples that can confound the analysis. By looking at the time points between 7 and 25 days, we sought to capture the kinetic variance in turion gene induction that may be exhibited by the two duckweed clones. Remarkably, many of these sorted genes with the highest expression at 25 days upon induction corresponded to genes identified as preferentially expressed in turions. Figure S12 shows the expression dynamics for some of these genes in the background of the two *S. polyrhiza* genotypes. Consistent with their high specificity for turions, we found that transcripts for these genes begin to show significant increase between 13- and 15-days post induction in the sp9512 background (Fig. S12A), while in sp9509 these genes only begin to show increase in their transcripts after 21 days (Fig. S12B). This variance in their induction kinetics mirrors the observed difference in turion induction timing between these two genotypes. For comparison to this group of turion associated genes that showed a clear delay of induction in the sp9509 genetic background, we also found genes that exhibited a markedly different behavior (Fig. S12C and S12D). OCT4 encodes an organic cation/carnitine transporter related to AtOCT4 and shows a similar response in both sp9512 and sp9509, with a peak of induction observed at 13 days post-treatment before increasing to high levels by 25 days. For the BI-1 cell death suppressor, we observed a peak of induction at 15 days before rising to high levels at 25 days in both genotypes. Finally, for PARP3, a seed-specific member of the conserved poly-(ADP-ribose) polymerase family, after a slight delay at the 15-day mark it was rapidly activated and reached much higher level of expression at the 21-day mark compared to the other 4 genes shown for the sp9509 genotype (compare Fig. S12B and Fig. S12D). The more rapid induction of these three genes in the sp9509 background in contrast to the process of turion formation indicates that they may also be induced in frond tissues in response to phosphate limitation, in addition to their later expression during turion biogenesis.

**Additional Materials and Methods**

**Starch analysis**

5 mg of dried *Spirodela polyrhiza* tissue were harvested from dry mass of normal fronds and turions samples. The samples were dissolved in 500 ml of 50mM sodium acetate pH 4.5 and ground (OPS Diagnostic, USA) 4000 rpm for 15 minutes. The ground mixture is then transferred to a glass tube and 1.5 ml of 50 mM sodium acetate, pH 4.5, is added and the solutions are autoclaved for 30 minutes to solubilize the starch. 500 ml of each autoclaved samples were incubated with 500 ml of 70 U/mg amyloglucosidase of *Aspergillus niger* diluted in 50mM sodium acetate pH 4.5 at 55°C for 30 minutes. The solutions were centrifuged at 14000 rpm for 5 minutes. The supernatants were measured using a YSI 2700 instrument (YSI, USA)

**Determination of chlorophyll content**

50 mg of fronds and mature turions from sp9512 and sp9509 were collected and transferred to an empty beating tube filled with one 4 mm silica bead. 1 mL of 80% (v/v) acetone was added to each tube and bead beating for 10 minutes at 4,000 RPM using a bead beater HT 6 OPS Diagnostic (Lebanon, NJ, USA) were performed to completely break the cells. The samples were transferred to a new 1.5 mL Eppendorf tube and incubated in dark at 4°C overnight. After incubation, samples are centrifuged at 16,000 g for 10 minutes at 4°C. The chlorophyll content was determined according to Arnon's method (Arnon, 1949).

**De Novo genome sequencing and assembly for *S. polyrhiza* 9512**

DNA was extracted from frozen tissue per previous methods with modifications (Lutz et al., 2011). Resulting DNA was quality controlled using Qubit for quantitation and gel electrophoresis to confirm the presence of high molecular weight (HMW) DNA. Samples with sufficient concentration and of HMW were used for sequencing. Unsheared HMW DNA was used to make Oxford Nanopore Technologies (ONT) ligation-based libraries. Libraries were prepared starting with 1.5 ug of DNA and following all other steps in ONT's SQK-LSK109 protocol. Final libraries were loaded on an ONT flowcell (v9.4.1) and run on the GridION. Bases were called in real-time on the GridION using the flip-flop version of Guppy (v3.1). The resulting fastq files were concatenated and used for downstream genome assembly steps. Illumina 2x150 paired-end reads were also generated for genome size estimates and polishing genome sequences. Libraries were prepared from HMW DNA using NEBNext (NEB, Beverly, MA) and sequenced on the Illumina NovaSeq. The resulting raw sequence was only trimmed for adaptors, resulting in >60x coverage of the sp9512 genome.

Resulting fastq files passing QC were assembled using our previously described pipeline (Michael, 2018) with the modification that the initial assembly was generated using FlyE (Kolmogorov *et al.*, 2019). The resulting assembly graph (gfa) was visually inspected with Bandage (v0.8.1) (Wick *et al.*, 2015). Consensus was generated with three rounds of mapping the ONT reads back to the assembly with minimap2 followed by Racon (v1.3.1) (Vaser *et al.*, 2017), and the final assembly was polished iteratively three times (3) using 2x150 bp paired-end Illumina reads mapped using minimap2 (v2.17-r941) (>98% mapping) followed by pilon (v1.22) (Walker *et al.*, 2014). The resulting assembly was assessed for traditional genome statistics including assessing genome completeness with Benchmarking Universal Single-Copy Orthologs (BUSCO) (v3) liliopsida odb10 database (Table 1) (Simão *et al.*, 2015).

**Gene prediction and annotation**

The chromosome resolved sp9512v2 genome was annotated using a pipeline consisting of four major steps: repeat masking, transcript assembly, gene model prediction, and functional annotation. Repeats were identified using EDTA (v1.9.8) (Ou *et al.*, 2019) and these repeats were used for softmasking. ONT cDNA reads were aligned to the genomes using minimap2 and assembled into transcript models using Stringtie (v1.3.6). The softmasked genome and Stringtie models were then processed by Funannotate (v1.6) (https://github.com/nextgenusfs/funannotate) to produce gene models. The resulting gene models were renamed reflecting the chromosome and the linear position on the chromosome. Predicted proteins were then functionally annotated using Eggnog-mapper (v2) (Huerta-Cepas *et al.*, 2017).

**Genome comparisons and synteny analysis**

The sp9509 and sp7498 genomes were downloaded (spirodelagenome.org). The SynMap tool on CoGe (Grover *et al*., 2017) and McScan python version (https://github.com/tanghaibao/jcvi/wiki/MCscan) were utilized to generate whole genome synteny maps, identify syntenic orthologs, and generate figures. Mcscan parameters were modified to test the hypothesis that *S. polyrhiza* was more fractionated across different clones. Longer range syntenic connections were identified by allowing syntenic blocks to only require 4 orthologs (min_size=4) in a span of 100 genes (dist=100).

**Annotation and characterization of LEA and oleosin genes**

The sp9512 genome was scanned using HMMER v.3.1b2 (Eddy, 2011) and Pfam hidden Markov models (HMMs) for late embryogenesis abundant (LEA) and oleosin genes. The following Pfam HMM models were used: PF01277 (Oleosin), PF03760 (LEA_1), PF03168 (LEA_2), PF03242 (LEA_3), PF02987 (LEA_4), PF00477 (LEA_5), PF10714 (LEA_6), PF00257 (Dehydrin), and PF04927 (SMP). Neighbor-joining (NJ) phylogenetic trees were constructed for LEA and oleosin genes using MUSCLE v.3.8.1551 (Robert, 2004) NJ phylogenetic trees were visualized using ggtree (Yu, 2020).

**RNA extraction from frond and mature turion tissues**

For RNA isolation from fronds, Fresh fronds were collected and frozen immediately in liquid nitrogen and stored at -80°C until needed. Total RNA isolated using the Life Technologies™ *mir*Vana™ miRNA Isolation kit (Catalog Number: AM1560). The tube of frozen frond tissue from sp9512 was filled with silica beads (0.5 g 100 µm silica beads, 0.5 g 1.7 mm zirconium beads, and 1 bead 4 mm silica bead). They were resuspended in 800 µL of the Lysis/Binding buffer. The tube was then frozen in liquid nitrogen for 30 seconds and bead beating for 10 minutes at 4000 rpm using a bead beater HT 6 OPS Diagnostic (Lebanon, NJ, USA) were performed to completely break the cells and release the nucleic acid. The supernatant was collected and transferred to a new 2 ml Eppendorf tube. To the new tube, 1/10 volume of miRNA Homogenate Additive is added and incubated on ice for 10 minutes. RNA extractions were performed according to Life Technologies™ *mir*Vana™ miRNA Isolation kit. RNA was eluted in 50 µL of preheated (95°C) DEPC-$H_2O$ directly onto the column membrane. It was then centrifuged at 10,000 RPM for 30 seconds at room temperature. RNA was transferred to a new 1.5 ml Eppendorf tube and stored at -80°C.

For RNA isolation from mature turions, fresh tissues from *S. polyrhiza* 9512 and 9509 were collected and frozen immediately in liquid nitrogen and stored at -80°C until needed. Total RNA was extracted using CTAB method (Gambino *et al*., 2008) with slight modifications to the protocol. Mature turions were ground into fine powder in liquid nitrogen. The powdered sample was transferred to bead beating tube that filled with silica beads (0.5 g 100 µm silica beads, 0.5 g 1.7 mm zirconium beads, and 1 bead 4 mm silica bead). They were resuspended in 800 µL of extraction buffer (2% CTAB, 2.5% PVP-40, 2 M NaCl,100 mM Tris-HCl pH 8.0, 25 mM EDTA pH 8.0 and 2% of β-mercaptoethanol added just before use). The tube was then frozen in liquid nitrogen for 30 seconds. After freezing the samples, bead beating for 15 minutes at 4000 rpm using a bead beater HT 6 OPS Diagnostic (Lebanon, NJ, USA). Thereafter, the supernatant was collected and transferred to a new 2 ml Eppendorf tube. The tube was incubated at 65°C for 10 min. After incubation, an equal volume of chloroform: isoamyl alcohol (24:1 v/v) was added to the tube and vortexed for 30-60 seconds and centrifuged at 10,000 RPM for 8 minutes at 4°C. The upper aqueous layer is carefully transferred into a new 2 ml Eppendorf tube. The supernatant was recovered and a second extraction with chloroform:isoamyl alcohol (24:1 v/v) was performed. The supernatant was transferred to new microcentrifuge tube and 100 microliter LiCl (3 M final concentration) was added. The mixture was incubated in -20°C for 30 min and RNA was selectively pelleted after centrifugation at 14,000 rpm for 30 min at 4°C. The pelleted was resuspended with 0.7 vols of cold isopropanol and immediately centrifuged at 14,000 rpm for 20 min at 4°C. The pellet was washed with ethanol (70%). This step is repeated for a total of 2 washes with ethanol (70%). The supernatant was centrifuged at 14,000 rpm for 20 min at 4°C. The pelleted was dried, resuspended in 50 µL DEPC–water and stored at -80°C. For the experiments using beads and mortar only, the isolations were carried out using similar buffer and procedure.

**RNA-seq library preparation and sequencing**

RNA samples were shipped and sequenced separately at either BGI Genomics (Hong Kong, China) or Novogene Co., Ltd (Beijing, China). For samples shipped to BGI Genomics, sequencing libraries were prepared using an in-house library prep kit. Briefly, mRNA was purified from total RNA using poly-T oligo-attached magnetic beads. First-strand cDNA was generated using random hexamer-primed reverse transcription followed by second-strand cDNA synthesis. Resulting cDNA was then subject to end repair, adenylation, and adaptor ligation. Fragments were purified using Ampure XP beads. Library quality was validated using an Agilent Technologies 2100 bioanalyzer. Libraries were then circularized, amplified with phi29, and paired-end sequencing (2 x 100bp) was performed using combinatorial Probe-Anchor Synthesis on the DNBSEQ G400 sequencing platform from MGI Tech Co., Ltd.

For samples shipped to Novogene, sequencing libraries were generated using the NEBNext Ultra RNA Library Prep Kit for Illumina (New England Biolabs, USA) following manufacturer's recommendations. mRNA was purified from total RNA using poly-T oligo-attached magnetic beads. First strand cDNA synthesis was performed using random hexamer primers followed by second-strand cDNA synthesis. Library fragments of 150-200 bp were purified with the AMPure XP system (Beckman Coulter, Beverly, USA) and amplified with Phusion High-Fidelity DNA polymerase. Library quality was determined with an Agilent Bioanalyzer 2100. Pair-end sequencing (2 x 150 bp) was performed on libraries using the Illumina Novaseq6000 S4 sequencing platform.

**RNA-Seq quality control and gene expression quantification**

To remove adaptor contamination and low quality-reads, clean reads were generated from all samples using fastp v.0.20.0 (Chen *et al.*, 2018) by filtering raw reads that contained adaptor contamination, more than 10 % uncertain nucleotides, or if low quality nucleotides (Q < 5) constituted more than 50 % of the read. This was performed with a modified version of fastp (https://github.com/novogene-europe/fastp) with the following parameters: --qualified_quality_phred 5, --unqualified_percent_limit 50, --n_base_limit 15, overlap_len_require 30, --overlap_diff_limit 1, --min_trim_length 10, --overlap_diff_percent_limit 10, -l 150, --json <library name>.fastp.json, --html <library

name>.fastp.html, --report_title <library name>.fastp.report. The high-quality cleaned reads (20,539,521 to 47,967,197) obtained from all constructed libraries were subjected to downstream analysis (Supplemental Table S1).

Clean reads were then mapped to the sp9512 genome using HISAT2 v.2.1.0 (Kim *et al.*, 2019) with the following parameter specified: --dta, --phred33. Alignment rate ranged from 96.65 to 97.56 % (Supplemental Table S1). Read counts were generated using feature Counts v.2.0.0[61] from the Subread software package by counting fragments (-p) that had both ends aligned to exons (-t exon) on the same chromosome (-B) on the same strand (-C). Successfully assigned alignments meeting these criteria ranged from 55.50 to 74.69 % (Supplementary Table S1).

**Gene expression level differences validated by qRT-PCR and PCR**

Total RNA was isolated from sp9512 mature turion and frond tissues using our current protocol. Then, the cDNA was synthesized using SuperScript III reverse transcriptase (Thermo Scientific, MA, USA). qRT-PCR was performed using the StepOnePlus Real-Time PCR System (Applied Biosystems, Thermo Fisher Scientific). The comparative threshold cycle method was used to determine relative gene expression, with the expression of SpActin as an internal control. PCRs were performed according to the protocol: initial denaturation at 95$^{o}$C for 10 min (stage 1), followed by 40 cycles (stage 2) of denaturation at 95$^{o}$C for 15 seconds, annealing at 60$^{o}$C for 1 min, and stage 3 (melt curve stage) at 95°C for 15 s, 60° C for 1 min and 95°C for 15 sec. Three technical replicates and three biological replicates were maintained for each sample. The relative expression levels were calculated as $2^{\Delta\Delta Ct}$. The primer sets for qRT-PCR are listed in Table S9.

For end point PCR, cDNA was synthesized from total RNA of sp9512 mature turion and frond tissues using SuperScript III reverse transcriptase (Thermo Scientific, MA, USA). PCR were conducted with the following parameter: initial denaturation at 94$^{o}$C for 1 min, followed by 35 cycles of denaturation at 94$^{o}$C for 45 seconds, annealing at 55$^{o}$C for 30 seconds, and extension at 72$^{o}$C for 5 minutes. The amplification products were analyzed by gel electrophoresis (Fig. S11). The primer sets for PCR are listed in Table S9.

**DNA extraction of frond and mature turion for methylation analysis**

Fresh fronds and mature turions of sp9512 were collected and frozen immediately in liquid nitrogen and stored at -80°C until needed. DNA was extracted using Qiagen isolation kit with slight modifications to the protocol. Fronds and Mature turions were ground into fine powder in liquid nitrogen. The powdered sample was transferred to bead beating tube that filled with silica beads (0.5 g 100 µm silica beads, 0.5 g 1.7 mm zirconium beads, and 1 bead 4 mm silica bead). They were resuspended in 800 µL of lysis buffer. The tube was then frozen in liquid nitrogen for 30 seconds. After freezing the samples, bead beating for 15 minutes at 4000 rpm using a bead beater HT 6 OPS Diagnostic (Lebanon, NJ, USA). Thereafter, the supernatant was collected and transferred to a new 2 ml Eppendorf tube. The tube was incubated at 65°C for 10 min. DNA extractions were performed according to Qiagen DNA Isolation kit. DNA was eluted in 50 µL of preheated (65°C) $H_2O$ directly onto the column membrane. It was then centrifuged at 10,000 RPM for 30 seconds at room temperature. DNA was transferred to a new 1.5 ml Eppendorf tube and stored at -80°C.

**DNA methylation calling and analysis**

Genomic reads from the raw nanopore fast5s generated from the frond and the turion samples were used for methylation calling as well as base called fastqs and the genome reference as described by Nanopore for phased methylation calling with a custom script (https://gitlab.com/salk-tm/phased-methylation). This entails mapping fastq reads to the genome and performing variant calling and haplotyping with Longshot (Edge & Bansal, 2019). Phased read IDs were used to parse the total methylation calls from Megalodon (https://nanoporetech.github.io/megalodon/index.html). Megalodon uses the raw fast5's and a reference to find methylation and was performed with the latest rerio models as "--guppy-config res_dna_r941_prom_modbases_5mC_v001.cfg'' and with the vcf from Longshot with "--variant-filename". The phased read IDs were used to parse out methylation calls using "megalodon_extras aggregate run" and passing in the megalodon directory and "--read-ids-filename" for each phased read file. The resultant bed file containing methylation calls at each "C" position was further parsed with a custom script (https://gitlab.com/NolanHartwick/bio_utils) using "split_methyl_bed.py" and the reference genome to generate three methylation bed files with the CpG, ChG, and Chh contexts. Differentially Methylated Regions (DMR) were determined between the frond and turion for CpG methylation using Metilene (Jühling *et al*., 2016). DMRs were considered significant where $q < 0.01$ where q is the q-value determined by the Benjamini-Hochberg false discovery rate procedure. DMRs were determined near differentially regulated genes with "bedtools intersect" (Jühling *et al*., 2016; Quinlan & Hall, 2010) within 2kb upstream or downstream of the gene start and end site, respectively. Finally, bed files were converted to BigWig files for visualization in the Integrated Genome Viewer (IGV) and Deeptools (Ramirez *et al*., 2016) for correlation comparisons and plotting. For chromosome-level methylation plots, the chromosome was divided into 100-Kb bins. A single average DNA methylation value was calculated for each bin, and bootstrap-based confidence intervals were determined from the distribution of individual cytosine methylation levels within each bin. For plots of gene body and TE profiles, the locus body – start-to-stop codon for genes and first to last bp for TEs – was divided into 10 proportional bins based on locus length. Additionally, 1000 bp upstream and downstream of genes or 500 bp upstream and downstream of TEs were divided into 10 proportional bins. A single DNA methylation level value was calculated for each bin across all loci, and bootstrap-based confidence intervals were determined from the distribution over loci.

The number of replicates (n) and the statistical details for each experiment are indicated in corresponding figure legends. Microsoft Excel 2010 was used for graph preparation and statistical analyses. Additional graphs were generated using the R programming language v.4.1.2 with the ggplot2 v.3.3.5 package.

***Supp data***

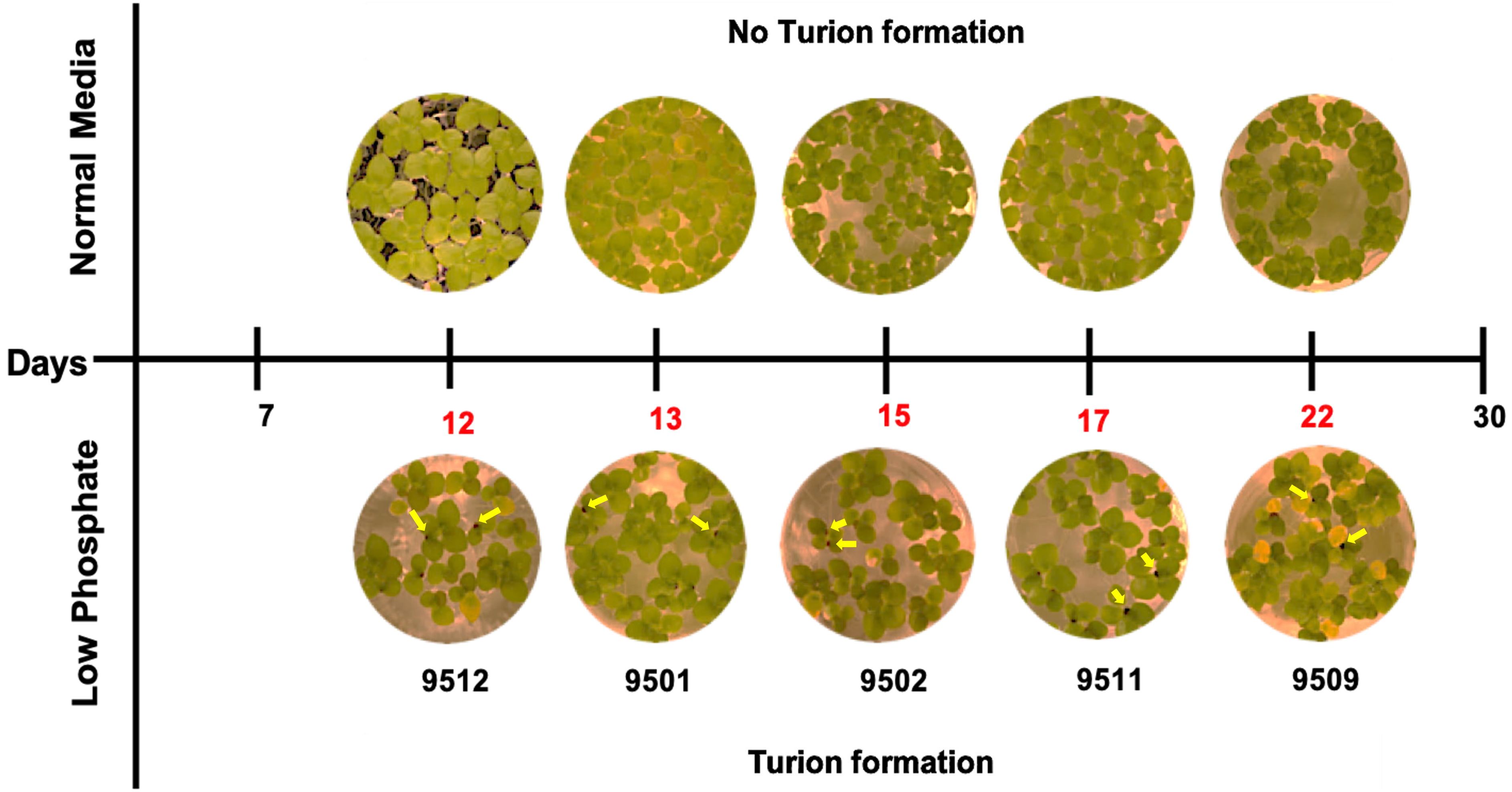


**Figure S1. Comparison of turion formation kinetics between five clones of *S. polyrhiza* with different STY**. Time of turion induction/commencement for each clone is highlighted in red numbers. Newly formed turions emerging from the meristem pockets are indicated by yellow arrows in the panels for induced fronds.

A. Physical parameter and chemical content in clones sp9512 vs. sp9509

| | Clone 9512 | | Clone 9509 | |
|---|---|---|---|---|
| | Fronds | Turions | Fronds | Turions |
| Chlorophyll (mg/g FW) | 0.51± 0.02 | 0.41 ± 0.02 | 0.65 ± 0.03 | 0.38 ± 0.02 |
| Starch (% DW) | 6.73 ± 1.40 | 75.06 ± 5.4 | 5.2 ± 1.07 | 79.71 ± 5.62 |
| Size of turions (diameter, mm) (n = 20) | ND | 0.99 ± 0.42 | ND | 1.57 ± 0.38 |
| Size (area, $mm^2$) (n = 20) | 31.5 ± 0.39 | ND | 32.69 ± 0.39 | ND |

Data presented as mean ± SD (n=3). ND = not detected, n = number of samples analyzed.

B. RNA yield and quality from sp9512 and sp9509 turions using MB method

| Samples | $A_{260/280}$ | $A_{260/230}$ | Concentration (ng/µl) |
|---|---|---|---|
| sp9512 | 1.74 | 2.50 | 60.8 |
| sp9509 | 1.84 | 2.30 | 90.2 |

C. RNA yield and quality isolated from sp9512 turions by different methods.

| Samples | $A_{260/280}$ | $A_{260/230}$ | Concentration (ng/µl) |
|---|---|---|---|
| sp9512- B | 1.55 | 1.20 | 35.2 |
| sp9512- M | 1.95 | 1.75 | 45.9 |
| sp9512- MB | 1.85 | 2.15 | 95.8 |

**Figure S2. Biochemical parameters comparison between turions and fronds for two *S. polyrhiza* clones**. (A) Physical parameters and chemical content in clones sp9512 vs sp9509. (B) Yields, as well as A260/280 and A260/230 ratios for total RNA isolated from sp9512 turions. (C) RNA yield and quality from turion of clones sp9512 vs sp9509. FW: fresh weight; DW: dry weight; B: beads only; M: mortar only; MB: beads and mortar with pestle

| | Sp9512 | sp9509 | sp7498 |
|---|---|---|---|
| Total length (bp) | 140,525,836 | 138,592,155 | 138,493,532 |
| Sequences (#) | 147 | 95 | 99 |
| Longest scaffold/Chr (bp) | 11,647,158 | 11,560,055 | 11,339,703 |
| Contig N50 length (bp) | 4,463,637 | 2,868,147 | 3,339,466 |
| Scaffold N50 length (bp) | 7,835,987 | 7,949,387 | 7,689,391 |
| Predicted genes (#) | 18,402 | 18,486 | 17,057 |
| Tandem repeat genes (#) | 2,751 | 2,814 | 2,156 |

**Figure S3*. Spirodela polyrhiza* 9512 de novo genome assembly statistics.**

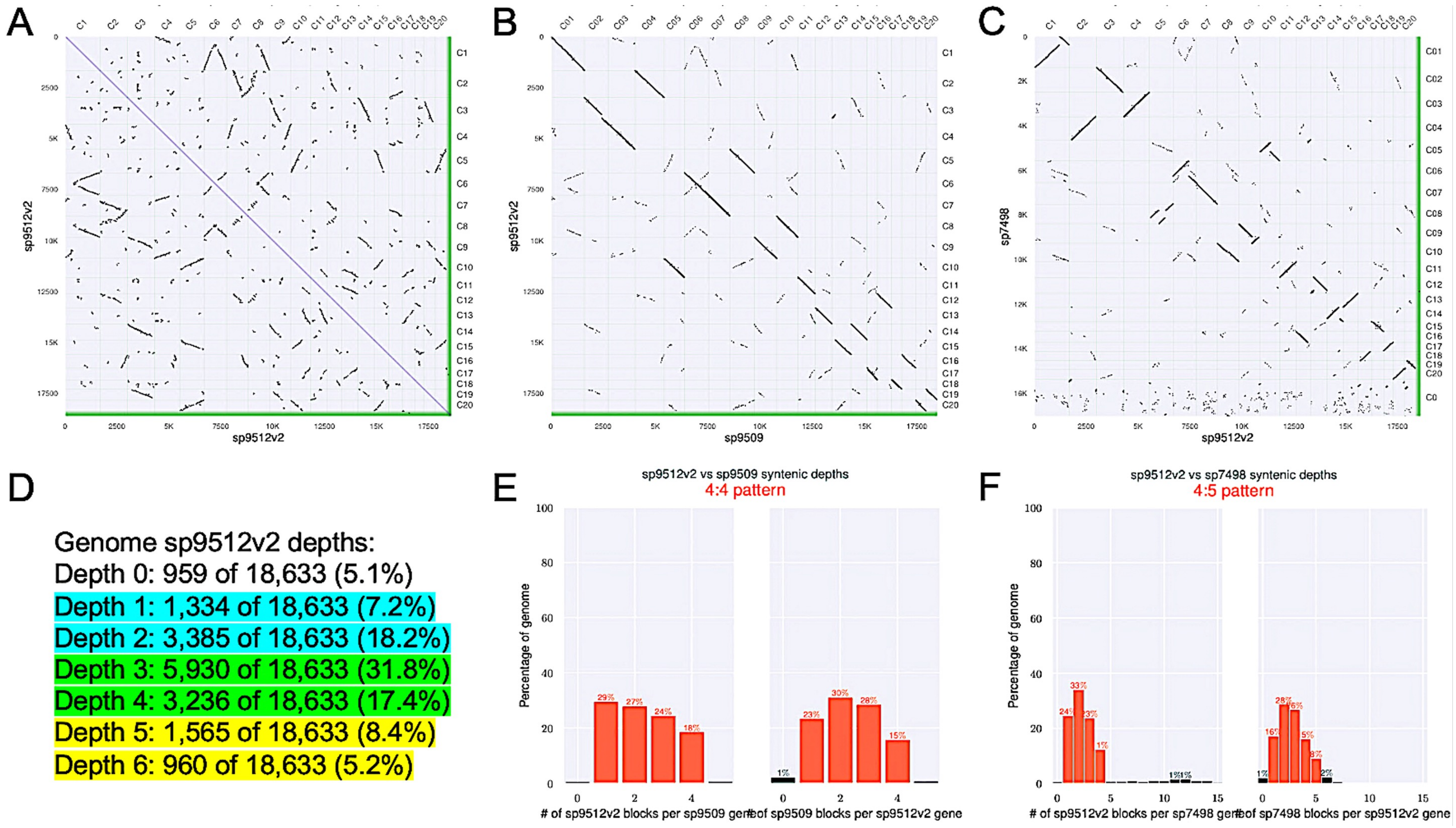


**Figure S4. Comparison of the sp9512 genome assembly with those from two published *Spirodela polyrhiza* reference genomes.** The sp9512v2 genome was compared to itself and the two high quality Spirodela reference genomes sp9509 and sp7498 for syntenic blocks defined by at least 4 proteins being syntenic within a span of 100 proteins. A) The dot plot comparing sp5912v2 against itself revealed the diagonal line representing the direct 1:1 self proteins, while the off diagonal are 4:4 remnants from the previous two whole genome duplications (WGD). B-C) Dot plot comparison of sp9512v2 to the reference genomes sp9509 and sp7498 revealed that the two genomes are highly co-linear without any large structural variations and evidence for the 4:4 retention of syntenic from the last two WGD. D) Summary of number and percent of proteins in syntenic blocks in the self-self sp9512v2 alignment. 5.1% of proteins were singletons and were not part of syntenic blocks. 7.2% and 18.2% of proteins were found in blocks one and two copies (highlighted in blue). 31.8% and 17% of proteins were found in blocks three and four copies (highlighted in green). 8.4% and 5.2% of proteins were found in blocks five and six copies (highlighted in yellow), which is consistent with Spirodela having two recent WGD events and the ancient tau WGD found in all monocots. E-F) Summary of syntenic blocks between sp9512v2 and sp9509 or sp7498 confirm the 4:4 ratio.

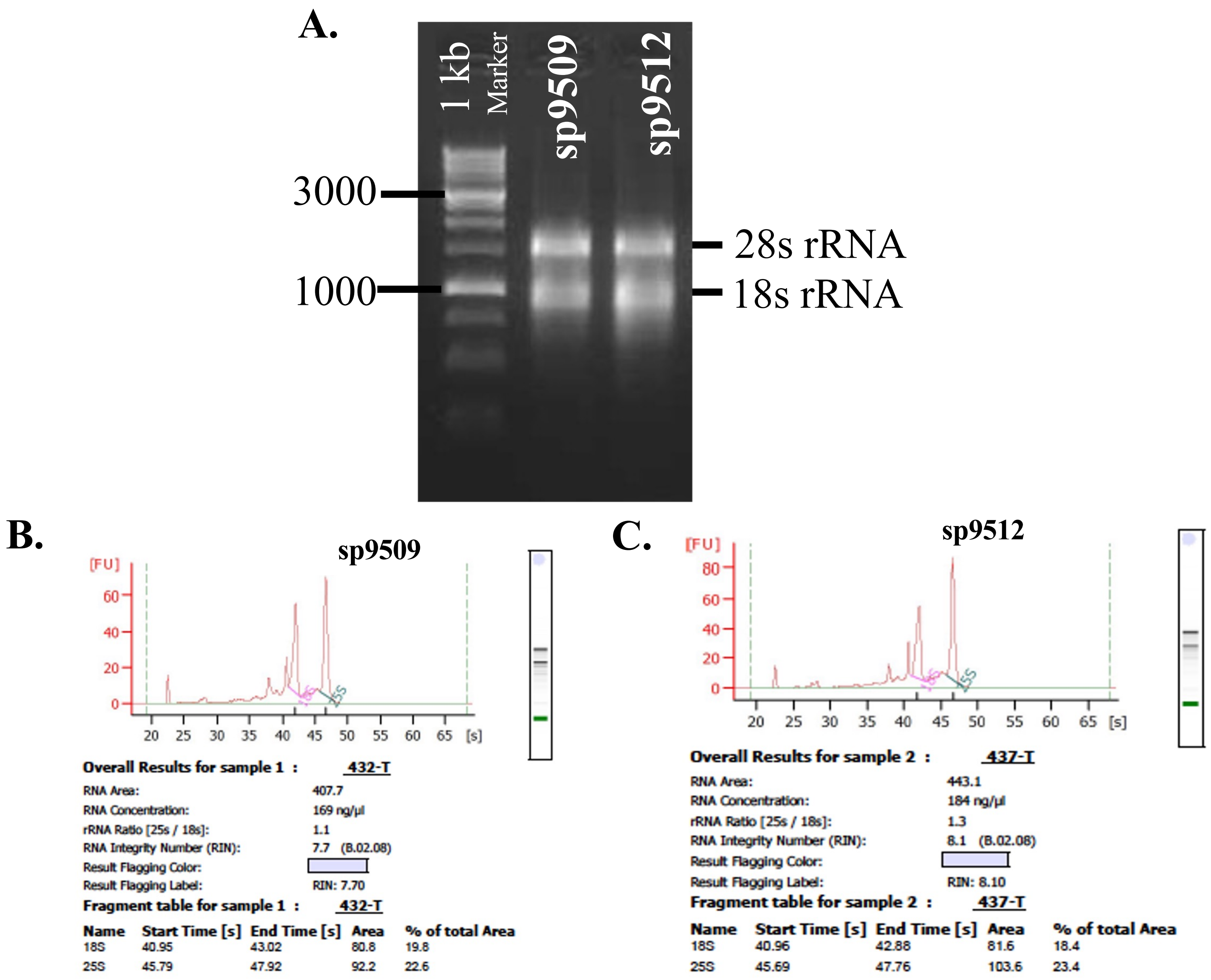


**Figure S5. Analysis of quantity and quality of Total RNA isolated from *S. polyrhiza* turions.** (A) Agarose gel electrophoresis of Total RNA isolated from *S. polyrhiza* turions. (B) The RNA quality of sp9509 detected by using Agilent 2100 Bioanalyzer. (C) The RNA quality of sp9512 detected by using Agilent 2100 Bioanalyzer. Electropherograms and gel–like images showed that total RNA isolated from turions are of good quality.

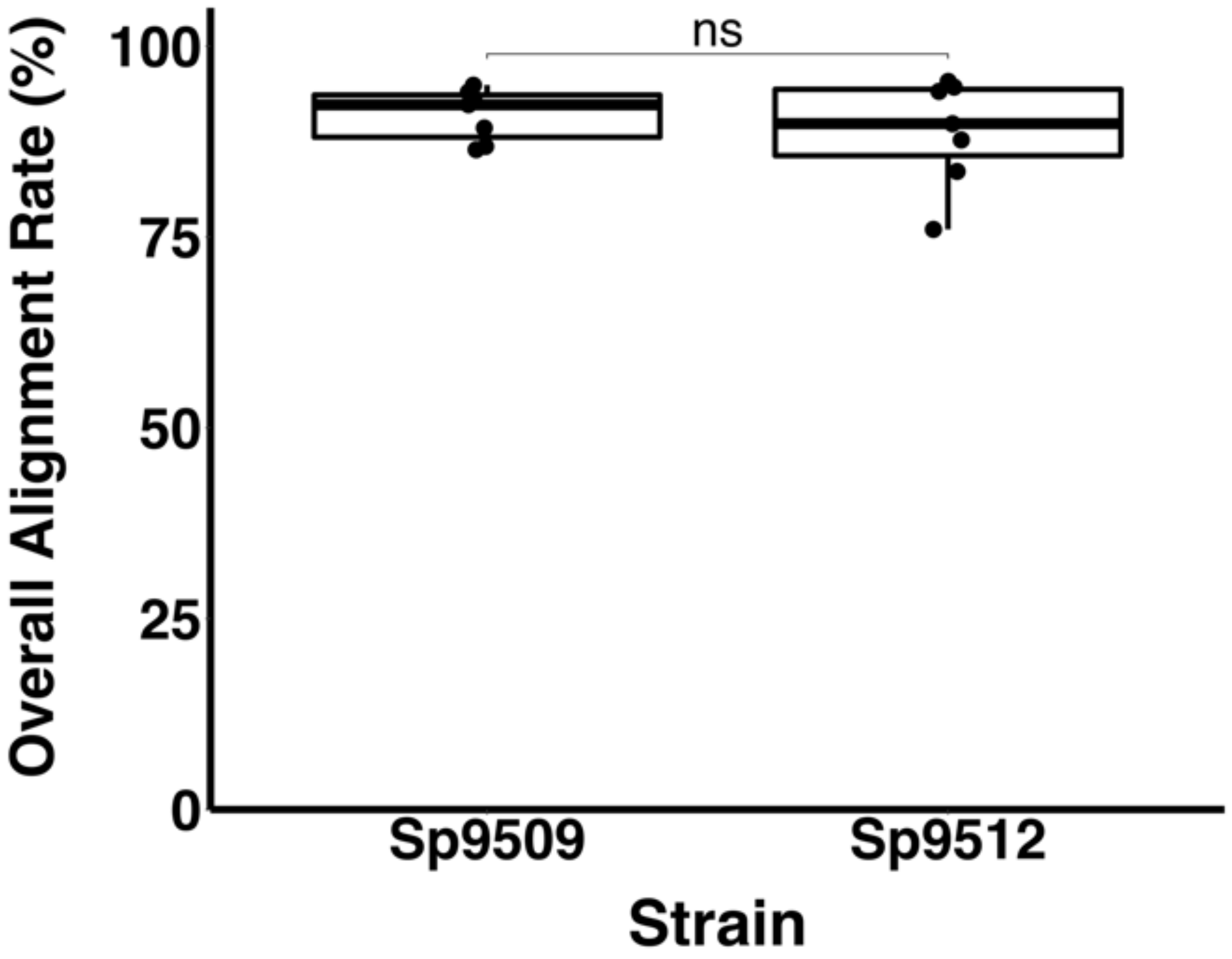


**Figure S6. Reference-based mapping of *S. polyrhiza* RNA-sequencing reads.** RNA-sequencing reads from all sample sets with either *S. polyrhiza* 9509 or *S. polyrhiza* 9512 were mapped to the *S. polyrhiza* 9509 reference genome using HISAT2. Similar results were obtained using the *S. polyrhiza* 9512 genome assembly as reference. Overall alignment rate was compared between strains using Welch's t-test with both reaching 90% or better. ns: not significant.

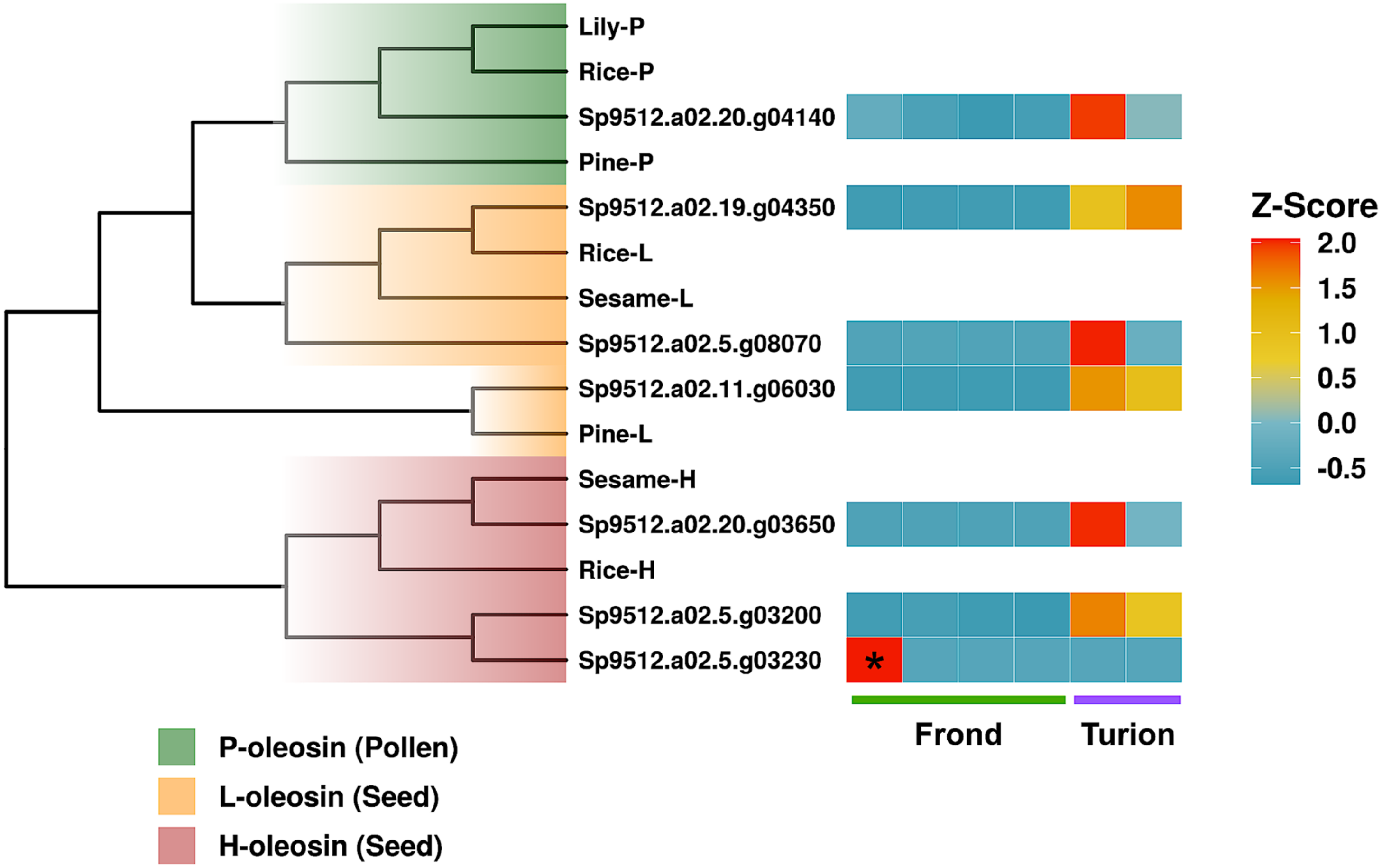


**Figure S7. Characterization of sp9512 oleosin gene members.** A hidden Markov model of the conserved hydrophobic domain from oleosins (PF01277) was used to retrieve oleosin proteins from sp9512 (66). A neighbor-joining phylogenetic tree is displayed comparing sp9512 oleosins to known oleosin proteins from sesame, rice, lily, and pine. Displayed is the relative gene expression for sp9512 oleosins for each sample compared to the mean expression (Z-score). Asterisks denote samples where gene read counts were less than 10. H-oleosin = high molecular weight oleosin (found in seeds); L-oleosin = low molecular weight oleosin (found in seeds); P-oleosin (found in pollen); Lily-P = ABK40507.1; Rice-P = BAF18317.1; Pine-P = AOZ15521.1; Rice-L = AAC02239.1; Sesame-L = AAD42942.1; Pine-L = AIC74542.1; Sesame-H = AAG23840.1; Rice-H = AAC02240.1

**Figure S8. Expression profile of sp9512 LEA genes.** Late-embryogenesis abundant (LEA) proteins in sp9512 were retrieved using hidden Markov models for 7 different LEA domains (45). LEA protein sequences were used to construct a neighbor-joining phylogenetic tree. Displayed is the relative expression of each LEA gene compared to the mean expression (Z-score) for sp9512 fronds and turions. sp9512 LEA proteins were also compared to LEA proteins from rye using BLASTP (45). Displayed are top rye hits using the top bit score. Rye LEA proteins expressed specifically in seed are colored in red.

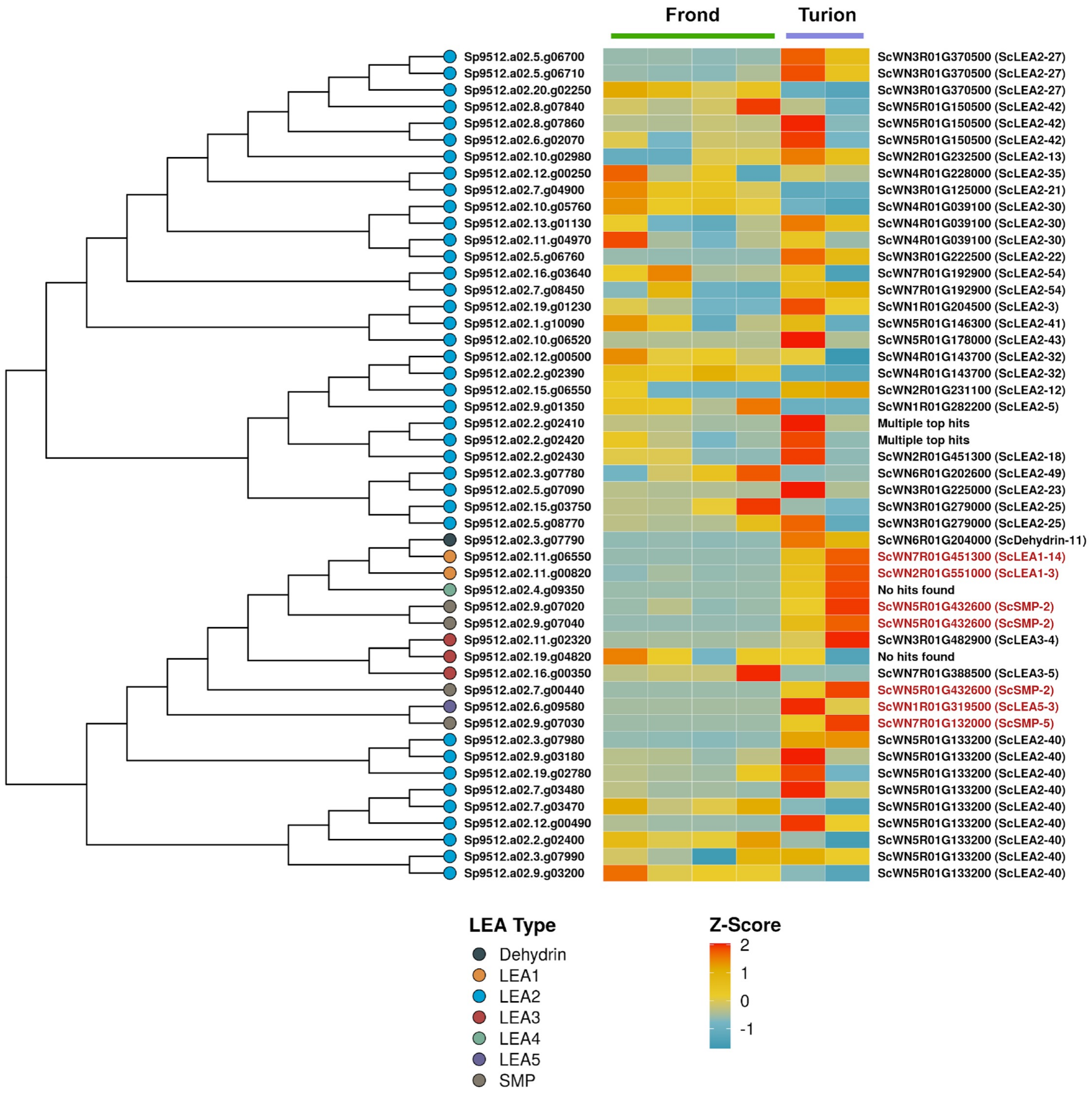

| Category | Term | Over_represented_pvalue | numDEInCat | numInCat | Ontology | Over_padj | Function |
|---|---|---|---|---|---|---|---|
| GO:0003777 | microtubule motor activity | 6.70E-16 | 17 | 42 | MF | 2.40E-12 | **Cell devision, DNA replication,** |
| GO:0005871 | kinesin complex | 6.70E-16 | 17 | 42 | CC | 2.40E-12 | **Cytoskeleton related** |
| GO:0003774 | cytoskeletal motor activity | 2.15E-15 | 17 | 46 | MF | 5.76E-12 | |
| GO:0005875 | microtubule associated complex | 1.17E-14 | 20 | 59 | CC | 2.50E-11 | |
| GO:0008574 | plus-end-directed microtubule motor activity | 3.13E-14 | 11 | 15 | MF | 5.60E-11 | |
| GO:0015630 | microtubule cytoskeleton | 9.76E-14 | 30 | 156 | CC | 1.50E-10 | |
| GO:0007018 | microtubule-based movement | 4.03E-13 | 18 | 58 | BP | 5.40E-10 | |
| GO:0007017 | microtubule-based process | 1.93E-11 | 26 | 135 | BP | 2.30E-08 | |
| GO:0005856 | cytoskeleton | 6.14E-11 | 33 | 227 | CC | 6.59E-08 | |
| GO:0006928 | movement of cell or subcellular component | 1.91E-09 | 19 | 91 | BP | 1.71E-06 | |
| GO:0007049 | cell cycle | 7.42E-09 | 33 | 280 | BP | 6.12E-06 | |
| GO:0022402 | cell cycle process | 4.77E-08 | 31 | 269 | BP | 3.66E-05 | |
| GO:0009524 | phragmoplast | 5.37E-08 | 12 | 44 | CC | 3.84E-05 | |
| GO:0000280 | nuclear division | 3.60E-07 | 18 | 120 | BP | 0.000241391 | |
| GO:0005819 | spindle | 1.16E-06 | 11 | 50 | CC | 0.000654285 | |
| GO:0006260 | DNA replication | 2.13E-06 | 14 | 86 | BP | 0.001144062 | |
| GO:0006268 | DNA unwinding involved in DNA replication | 3.88E-06 | 4 | 5 | BP | 0.001984001 | |
| GO:0000278 | mitotic cell cycle | 6.90E-06 | 20 | 168 | BP | 0.003084065 | |
| GO:0008017 | microtubule binding | 2.56E-05 | 10 | 58 | MF | 0.009158 | |
| GO:0140013 | meiotic nuclear division | 2.87E-05 | 11 | 67 | BP | 0.009929119 | |
| GO:0006261 | DNA-dependent DNA replication | 3.49E-05 | 11 | 73 | BP | 0.011357369 | |
| GO:1903046 | meiotic cell cycle process | 7.96E-05 | 13 | 96 | BP | 0.021919059 | |
| GO:1903047 | mitotic cell cycle process | 0.000139297 | 17 | 156 | BP | 0.03250174 | |
| GO:0051321 | meiotic cell cycle | 0.000149753 | 13 | 103 | BP | 0.034197877 | |
| GO:0000911 | cytokinesis by cell plate formation | 0.000163219 | 8 | 36 | BP | 0.034451367 | |
| GO:1902410 | mitotic cytokinetic process | 0.000163219 | 8 | 36 | BP | 0.034451367 | |
| GO:0015631 | tubulin binding | 0.00017904 | 10 | 72 | MF | 0.036257214 | |
| GO:0010564 | regulation of cell cycle process | 0.000190309 | 12 | 94 | BP | 0.037390403 | |
| GO:0032506 | cytokinetic process | 0.000191603 | 8 | 37 | BP | 0.037390403 | |
| GO:0048285 | organelle fission | 6.92E-07 | 20 | 152 | BP | 0.000412623 | **Developmental programs** |
| GO:0009647 | skotomorphogenesis | 0.000105177 | 6 | 14 | BP | 0.026877681 | |
| GO:0009640 | photomorphogenesis | 0.000134976 | 10 | 41 | BP | 0.032193256 | |
| GO:0015698 | inorganic anion transport | 2.11E-05 | 11 | 50 | BP | 0.008569148 | **Transporters** |
| GO:0008509 | anion transmembrane transporter activity | 2.25E-05 | 19 | 132 | MF | 0.008611363 | |
| GO:0022857 | transmembrane transporter activity | 2.56E-05 | 43 | 452 | MF | 0.009158 | |
| GO:0015103 | inorganic anion transmembrane transporter activity | 4.73E-05 | 10 | 44 | MF | 0.014410584 | |
| GO:0015075 | ion transmembrane transporter activity | 7.49E-05 | 30 | 292 | MF | 0.021146011 | |
| GO:0098656 | anion transmembrane transport | 8.50E-05 | 19 | 143 | BP | 0.022801542 | |
| GO:0015112 | nitrate transmembrane transporter activity | 0.000100051 | 5 | 9 | MF | 0.026191497 | |
| GO:0005215 | transporter activity | 0.000131811 | 44 | 519 | MF | 0.03215283 | |
| GO:0006820 | anion transport | 0.000159425 | 21 | 176 | BP | 0.034451367 | |
| GO:0034220 | ion transmembrane transport | 0.000228861 | 32 | 330 | BP | 0.043094151 | |
| GO:0015706 | nitrate transport | 0.000267057 | 5 | 11 | BP | 0.048581674 | |
| GO:0010345 | suberin biosynthetic process | 0.000168431 | 7 | 20 | BP | 0.034764717 | **Cell wall metabolism** |
| GO:0005618 | cell wall | 0.000278767 | 33 | 305 | CC | 0.049049242 | |
| GO:0030312 | external encapsulating structure | 0.000278767 | 33 | 305 | CC | 0.049049242 | |

**Figure S9.** Overrepresented Gene Ontology (GO) terms for up-regulated genes in fronds compared to turions. A Log2FC of 3 was used as a cutoff for inclusion of the DEGs for this analysis.

| Seq ID | Normalized counts | | | | | | Gene description | Function |
|---|---|---|---|---|---|---|---|---|
| | Frond | | | | Turion | | | |
| sp9512.a02.5.g05700 | 73 | 66 | 63 | 33 | 6322 | 8127 | GDSL esterase/lipase | Lipid metabolism |
| sp9512.a02.9.g07030 | 0 | 0 | 0 | 0 | 1220 | 3418 | Seed maturation protein | Seed development |
| sp9512.a02.11.g07860 | 0 | 0 | 0 | 1 | 20 | 20 | Homeobox-leucine zipper protein HAT22 | Transcription factor |

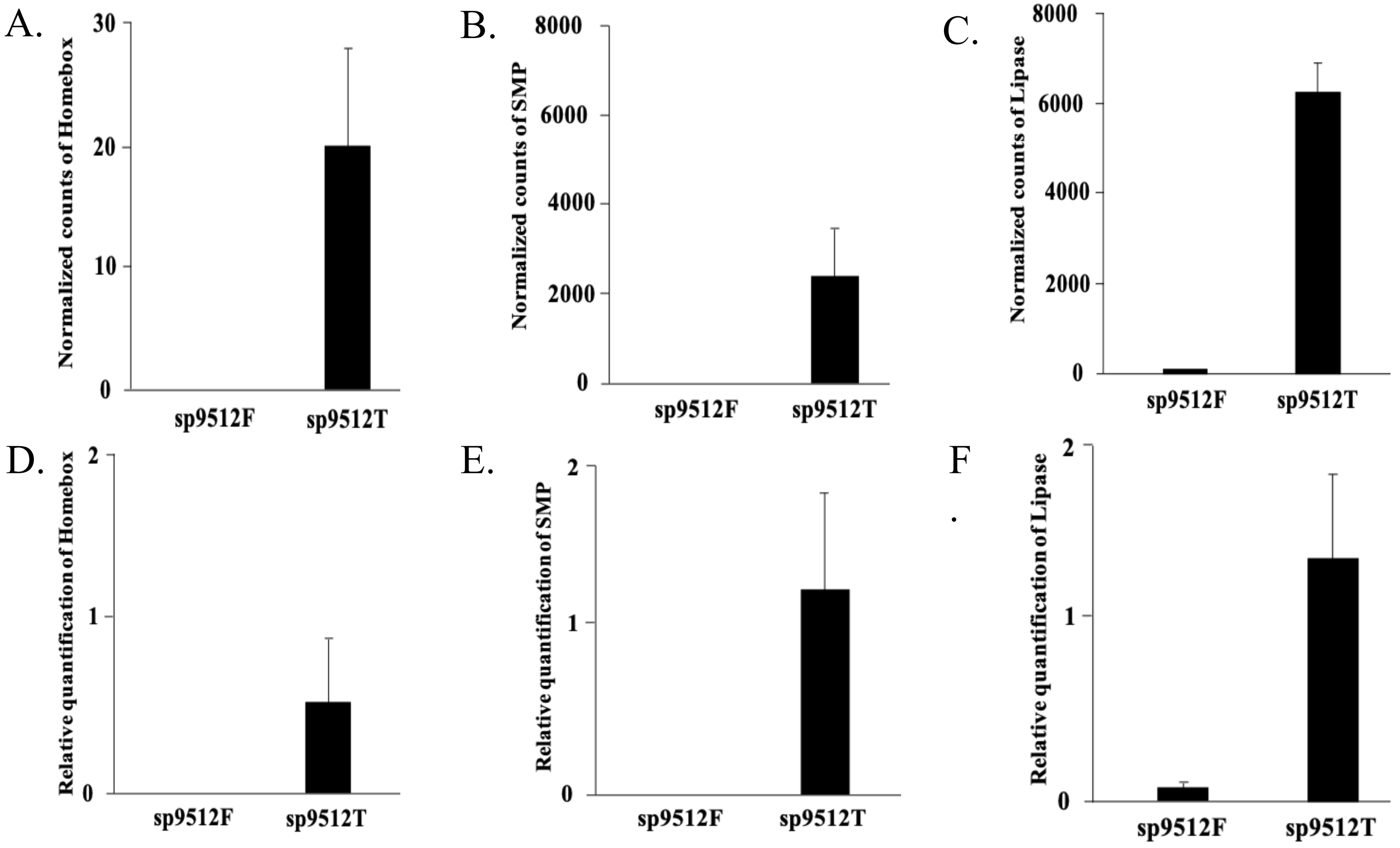


**Figure S10. Gene expression and qRT-PCR verification of three turion-associated genes.** (A-C) Graphs showing the average gene expression in the transcriptome dataset (DESeq2 normalized counts) of three sp9512 turion-associated genes (D-F) RT-qPCR verification of three turion associated genes in sp9512. sp9512F = sp9512 fronds, sp9512T = sp9512 turions. Homeobox gene ID: sp9512.a02.11.g07860, SMP (Seed Maturation Protein) gene ID: sp9512.a02.9.g07030, GDSL esterase/lipase gene ID : sp9512.a02.5.g05700, respectively.

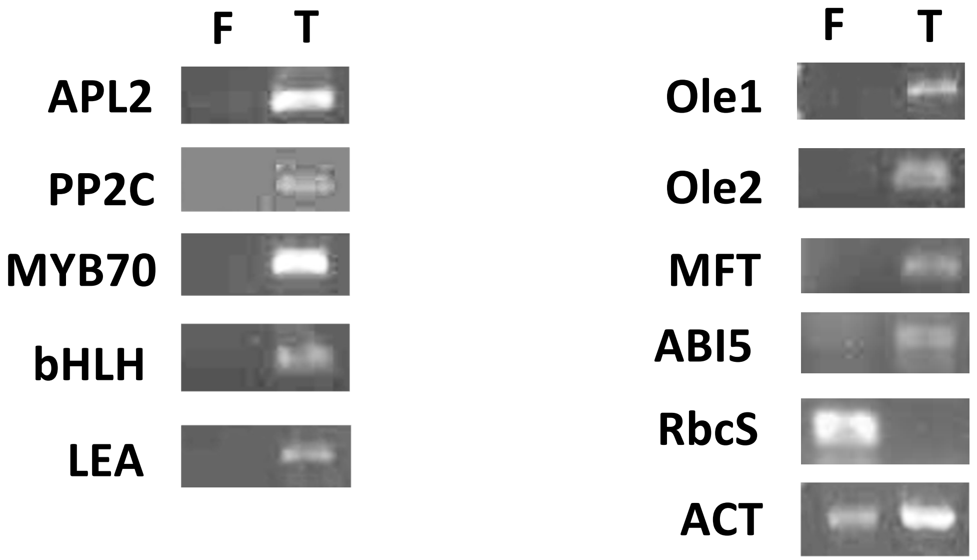


**Figure S11. sp9512 Turion Genes validation via RT-PCR**. Selection of overrepresented genes in Turion sp9512 were validated using end-point PCR. APL2 gene ID: sp9512.a02.20.g05195, PP2C gene id: sp9512.a02.15.g02700, MYB70 gene id:sp9512.a02.10.g05540, BHLH gene id:sp9512.a02.13.g07090, LEA gene ID : sp9512.a02.7.g00440, Ole1 gene ID: sp9512.a02.19.g04350, Ole2 gene ID: sp9512.a02.11.g06030, MFT gene id: sp9512.a02.5.g02270_p1, ABI-5 gene id: sp9512.a02.8.g03980_p1, RbcS gene id: Sp9512.a02.5.g01620_p1, ACT gene ID: sp9512.a02.6.g00520, respectively. F: Frond, T: Turion, APL2: AGPase Large subunit 2, PP2C: Protein phosphatase 2C, MYB70: , bHLH: Basic Helix Loop, Ole : Oleosin, LEA : late embryogenesis, MFT: MOTHER of FT and TF, ABI5: ABSCISIC ACID-INSENSITIVE 5, RbcS: RuBisCO, ACT : actin.

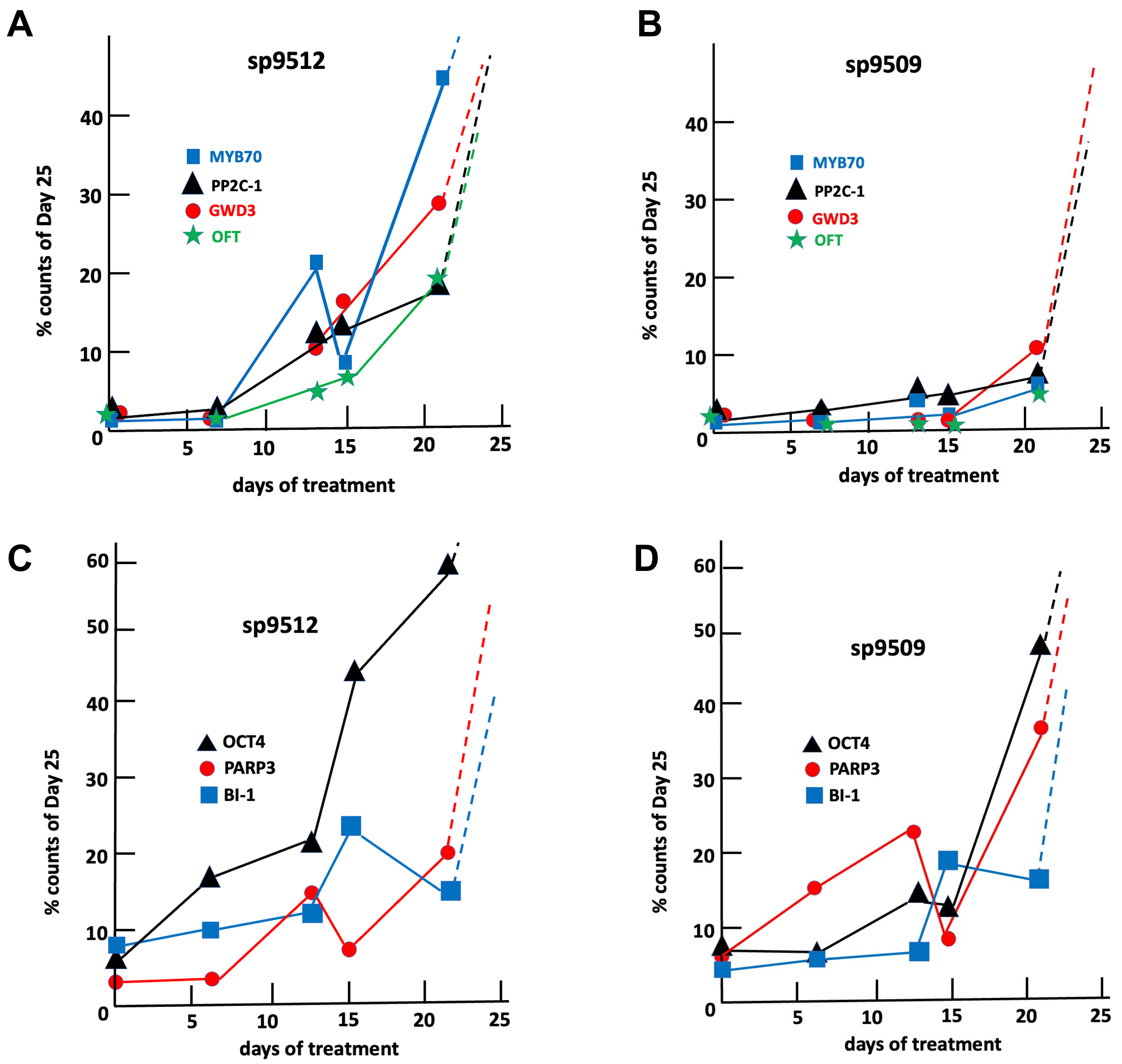


**Figure S12. Comparative transcriptomics identified transcripts with distinct behaviors between two *S. polyrhiza* genotypes.** Transcriptome normalized reads from sp9512 and sp9505 were mapped to the annotation of the sp9512v2 genome assembly. Reads from two to four replicates were averages at each time point and the reads at 25 days for each set of data were set as the 100% level in order to compare the different genes. Reads from fronds grown for 7 days under normal media were used as the zero-time control for reads from samples collected at 7, 13, 15, 21 and 25 days of growth in media with 2 uM phosphate. **Panels A and C**: reads from sp9512 frond samples. **Panels B and D**: reads from sp9509 samples. Gene numbers: MYB70 (Sp9512.a02.10.g05540_p1), PP2C-1 (Sp9512.a02.15.g02700_p1), GWD3 (Sp9512.a02.6.g00030_p1), OFT (Sp9512.a02.4.g00550_p1), OCT4 (Sp9512.a02.18.g03490_p1), PARP3 (Sp9512.a02.2.g00580_p1) and BI-1 (Sp9512.a02.11.g03070_p1).

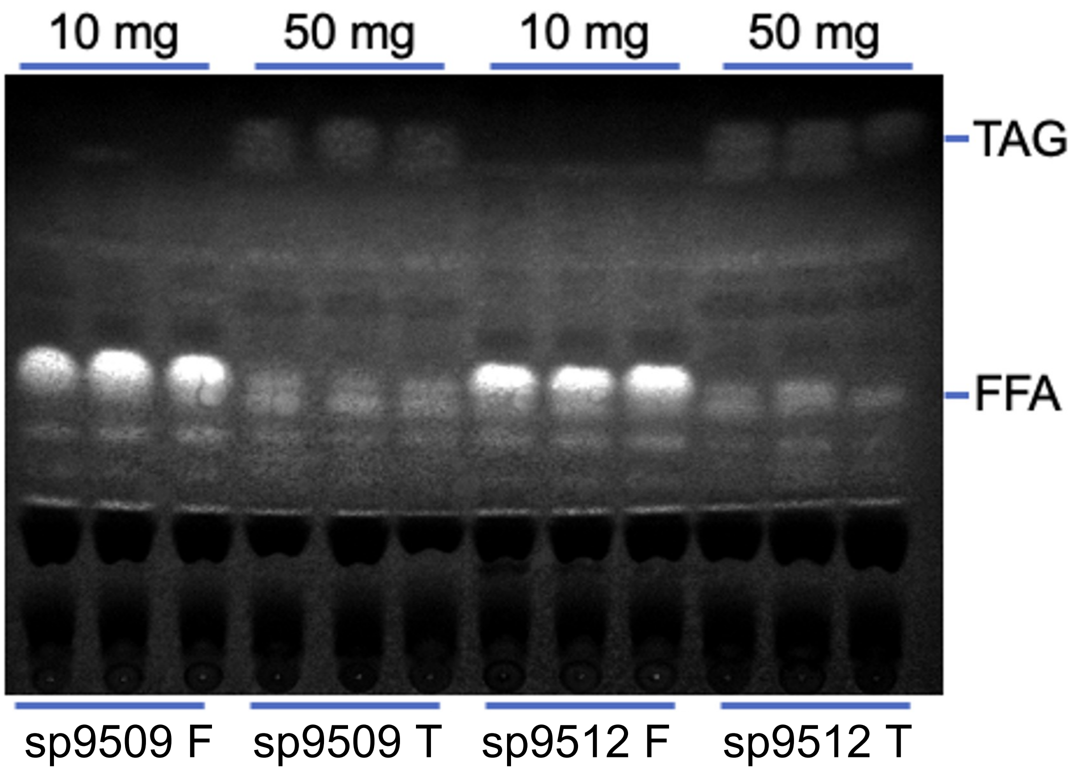


**Figure S13. Analysis of triacylglycerol (TAG) and free fatty acid (FFA) in *Spirodela polyrhiza* tissues**. Fronds (F) and turions (T) from sp9509 and sp9512 were freeze-dried before extraction with solvents. The extracts are separated on a TLC plate and stained with primuline to reveal the fatty acids. The total amount of tissue (dry weight) used to make extracts for analysis are shown on top for each set of samples. Each sample was analyzed in triplicates.

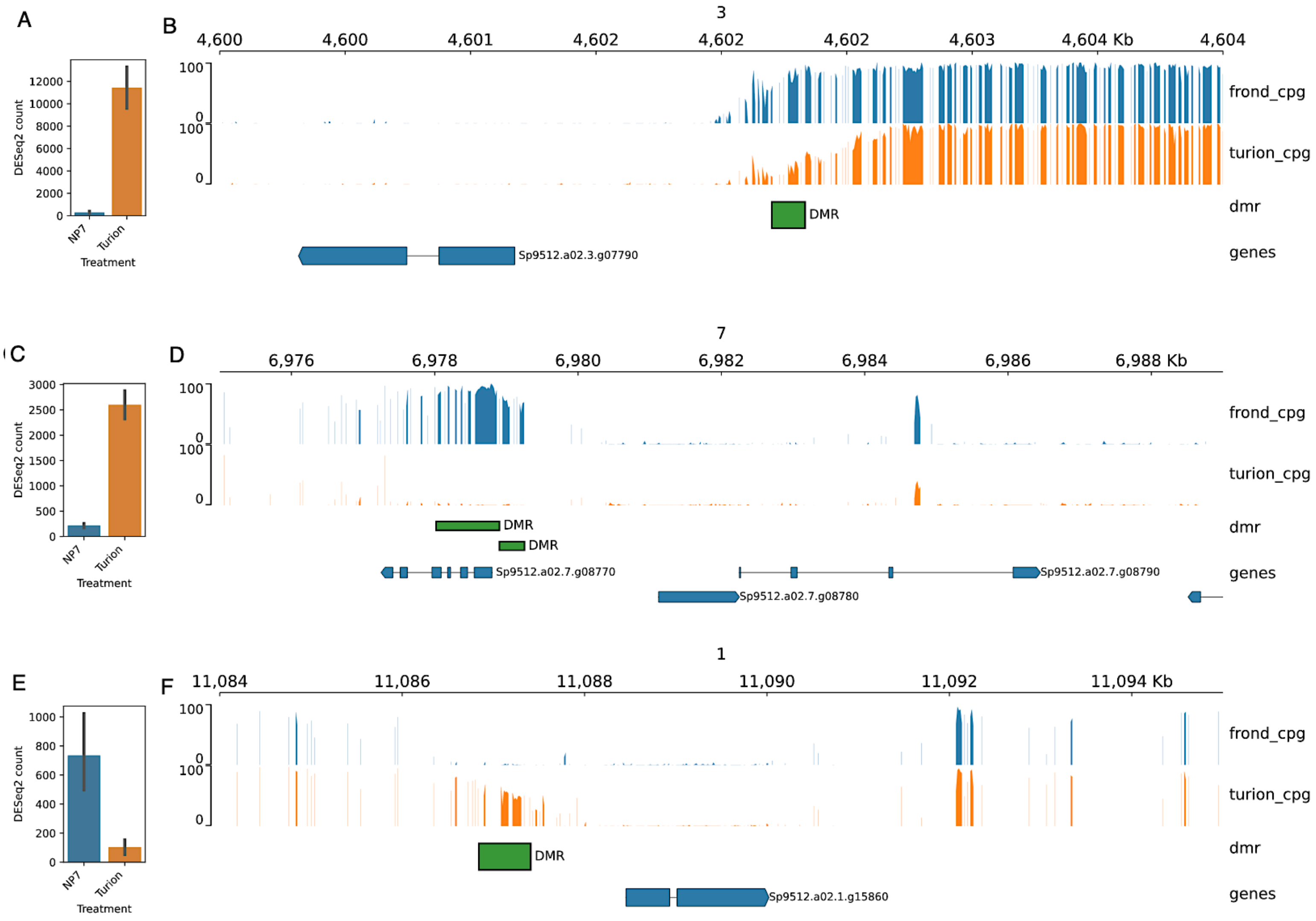


**Figure S14. Candidate loci in sp9512 that may be under epigenetic control via differential CpG methylation.** DMRs were mapped to regions of the sp9512 genome that exhibit either decreased (panels B and D) or increased (panel F) CpG methylation in turion gDNA vs those of fronds. The genes closest to these DMRs in these cases exhibited also the reciprocal changes in their transcript levels between the two tissues, with heightened expression observed for lower CpG methylation (panels A and C) while the opposite is observed in the latter case (panel E). Sp9512.a02.3.g07790_p1 (panel B) encodes a possible dehydrin class of LEA; Sp9512.a02.7.g08780_p1 (panel D) encodes a RING-H2 finger protein; Sp9512.a02.1.g15860_p1 (panel F) likely encodes a UDP-glycosyltransferase.

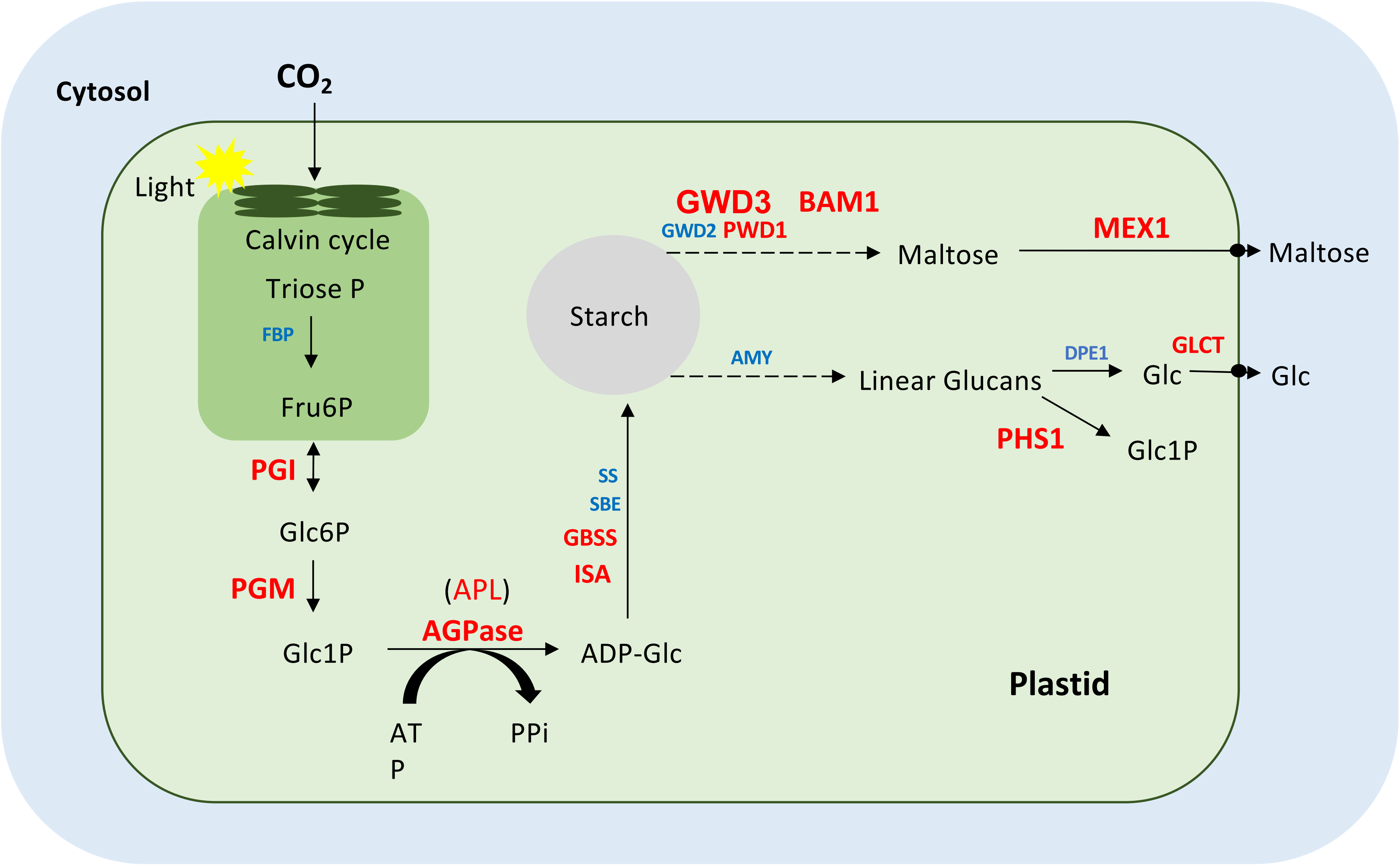


**Figure S15. Starch biosynthesis and degradation pathways an associated genes in sp9512**. Transcripts of mature turions as compared to fronds that are identified in the pathways are labeled in blue (down regulated) and red (up regulated). FBP: Fructose-Bisphosphatase 1, PGI: Glucose-6-phosphate isomerase, PGM: Phosphoglucomutase, AGPase: ADP-glucose pyrophosphorylase, ISA: Isoamylase, GBSS: Granule-bound starch synthase, SBE: starch-branching enzyme, SS: starch synthase, GWD: $\alpha$-Glucan water dikinase, PWD: Phosphoglucan water dikinase, BAM: Beta amylase, AMY: Alpha amylase, MEX1: Maltose excess protein 1, DPE1: 4-alpha-glucanotransferase DPE1, PHS1: phosphorylase isozyme, GLCT: glucose transporter, Glc: glucose.

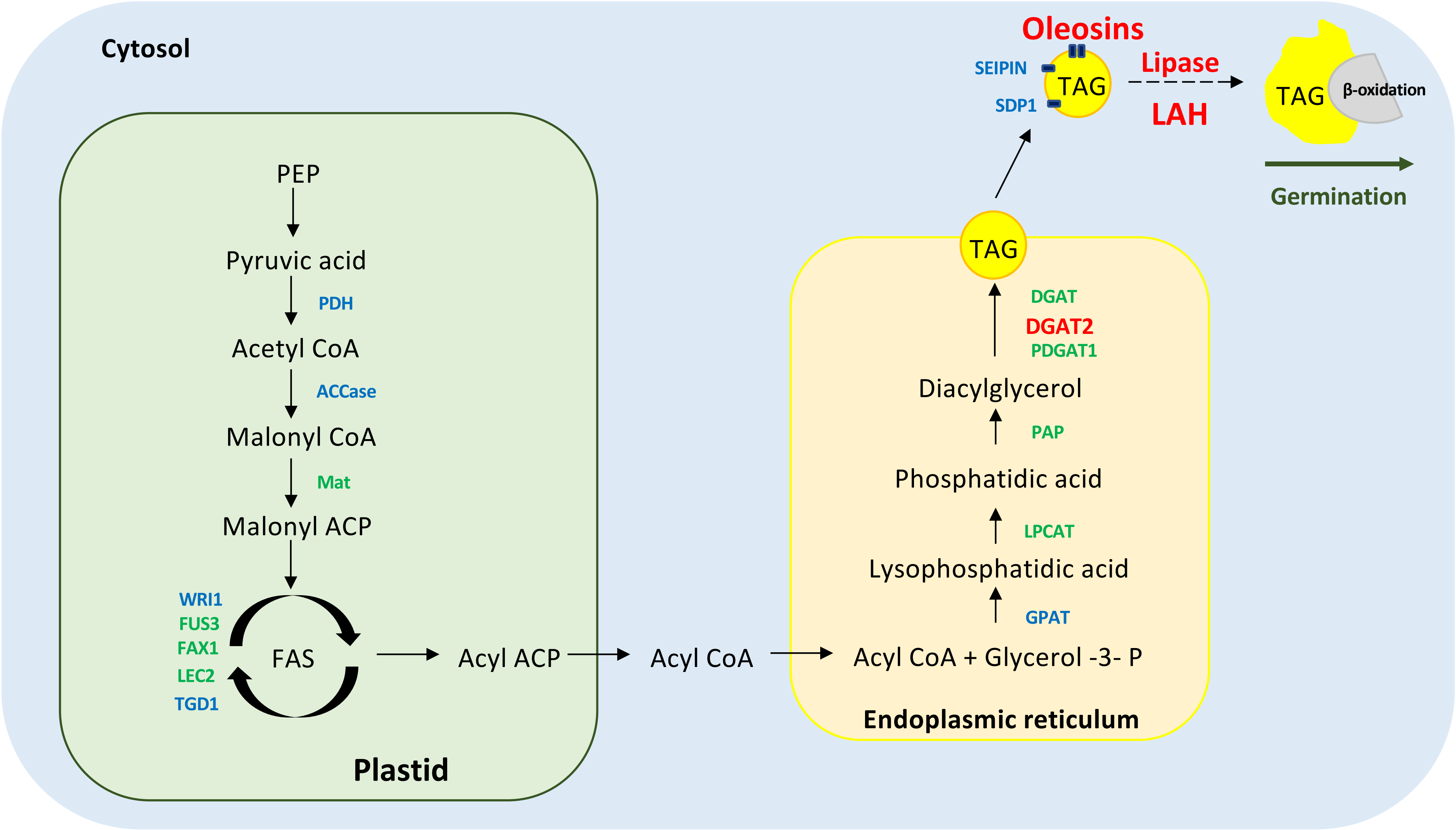


**Figure S16. TAG biosynthesis and degradation pathways and genes of sp9512.** Genes with transcripts identified in mature turions and their relative read counts to those of fronds are labeled in blue (down regulated), green (no significant difference), and red (up regulated). PDH: Pyruvate dehydrogenase, ACCase: Acetyl-CoA carboxylase, Mat: Methionine adenosyltransferase, FAS: Fatty acid synthase, WRI1: WRINKLED1, FUS3: FUSCA3, FAX1: Fatty acid export 1, LEC2: Leafy Cotyledon 2, TGD1: Trygalactosydiacylglycerol 1, GPAT: Glycerol-3-phosphate acyltransferase, LPCAT: Lysophosphatidylcholine acyltransferase, PAP: phosphatidic acid phosphatase, DGAT: diacylglycerol acyltransferase, SDP1: sugar-dependent 1, LAH: Lipid acyl hydrolase.
